# Vancomycin promotes key microbiota-pathogen interactions and removes protective bottlenecks to enteric infection

**DOI:** 10.64898/2026.06.14.730666

**Authors:** Sarah E. Woodward, Jorge Peña-Díaz, Antonio Serapio-Palacios, Stefanie L. Vogt, Madeline A. Wang, Wenny Feng, Kelsey E. Huus, Zakhar Krekhno, Laurel M.P. Neufeld, James C. Forward, Mihai Cirstea, B. Brett Finlay

## Abstract

Antibiotic exposure disrupts enteric pathogen colonization resistance, yet how antibiotics reshape pathogen population dynamics, infection bottlenecks, and strain-level heterogeneity in the gut remains poorly understood. Here, we combine high-resolution pathogen barcoding, transcriptomic, and metabolomic analyses to quantify how short-term vancomycin perturbation alters infection ecology *in vivo*. We use *Citrobacter rodentium* as a model for human infection by pathogenic *Escherichia coli*–an antimicrobial resistance priority group–to demonstrate that just two days of vancomycin pre-treatment profoundly reshapes infection trajectories, driving rapid, global gut colonization, a dramatic increase in pathogen founding population size, and preservation of strain diversity across intestinal sites. Notably, vancomycin eliminated the hallmark heterogeneity of *C. rodentium* infection, resulting in fully reproducible colonization across hosts. Population-level analysis revealed that antibiotic treatment relaxes competitive constraints both with the resident microbiota and among clonal pathogen lineages, allowing early-established founders to persist and expand. Despite accelerated pathogen engraftment and tissue pathology, transcriptomic analysis revealed reduced virulence gene expression. Instead, antibiotic-induced metabolic restructuring of the gut created permissive conditions for pathogen expansion. Interactions with a vancomycin-altered microbiota, dominated by *Akkermansia* and *Bacteroides*, further promoted nutrient cross-feeding and influenced epithelial attachment. Together, we illustrate how short-term antibiotic exposure reshapes enteric infection by removing ecological bottlenecks that normally constrain strain diversity and infection outcomes. These findings have implications for antibiotic use, antimicrobial resistance transmission, and therapeutic strategies that rely on competition-driven dynamics, such as strain replacement.

## Introduction

A recent analysis of global antibiotic use from 2000-2021 estimated that, on average, 30% of antibiotics were used inappropriately^1^. In Canada, over the course of 2019, antibiotics were prescribed at a rate of 627.3 prescriptions per 1000 people, of which more than half were estimated to be inappropriate or unnecessary^2^. Antibiotics are continually overused despite long-known disease associations and their status as a modifiable risk factor. Antibiotics have both direct and indirect effects on the host environment that may impact pathogen colonization. For example, bactericidal antibiotics such as ciprofloxacin, ampicillin, and kanamycin have been shown to induce reactive oxygen production in host cells, leading to mitochondrial dysfunction and affecting the integrity of the epithelial barrier^3^. The vancomycin-associated microbiota has been shown to alter the host immune response, skewing human CD4^+^ T cells towards a Th1/Th17 response and producing higher levels of TNF^4,5^.

Antibiotic use is a known driver of microbial dysbiosis and disruption of colonization resistance to infection. One study of hospital-acquired *Clostridioides difficile* (formerly *Clostridium difficile)* infection across Europe identified that 92% of positive patients reported antibiotic use within 3 months of hospital admission, suggesting that antibiotic use may increase susceptibility to infection^6,7^. Despite the association between antibiotic use and pathogen infection, further antibiotic administration remains the treatment of choice for infection with *C. difficile* and most other enteric pathogens. Vancomycin, which targets Gram-positive bacteria by inhibiting cell wall synthesis, is recommended for the treatment of severe *C. difficile* infection and methicillin-resistant *Staphylococcus aureus* (MRSA)^8,9^. Host exposure to vancomycin has been shown to increase susceptibility to the enteric pathogen *Salmonella enterica* serovar Typhimurium^10^, attributed to changes in the gut metabolomic profile, particularly alterations in bile acid, steroid, and tryptophan concentrations^11^.

The rise of antimicrobial resistance (AMR) is a major consequence of compounding antibiotic overuse which threatens our ability to treat bacterial infections, such as those that cause diarrheal disease. The World Health Organization (WHO) has identified several AMR priority pathogens, many of which are causative agents of diarrheal disease. Every year more than 500,000 children die of diarrheal diseases, predominantly in low– and middle– income countries. Pathogenic *Escherichia coli* are a leading cause of death in children under age five, with rising incidence of AMR, making them one of the top-ranked WHO priority pathogens^12,13^. Furthermore, antibiotic treatment removes critical colonization resistance from the resident microbiota, leaving hosts vulnerable to subsequent *E. coli* colonization and disease. Studies in mice have identified that antibiotic treatment before challenge with *Citrobacter rodentium*, a model of human pathogenic *E. coli* infection, increases disease severity in an antibiotic-dependent manner^14^. This effect is largely shown in response to antibiotic pre-treatments targeting Gram-negative bacteria, consistent with direct competition with Gram-negative *E. coli*. However, off-target effects of antibiotics targeting Gram-positive bacteria may also impact *E. coli* colonization and pathogenesis.

*C. rodentium* shares many of the virulence factors and disease characteristics which define human infection by Enterohemorrhagic *E. coli* (EHEC). These include the hallmark Type III Secretion System (T3SS) which resembles a molecular syringe that temporally injects protein effectors directly into the host epithelium to manipulate host cell processes and cause disease. The T3SS is essential for *C. rodentium* (and EHEC) colonization and virulence. Regulation of the system, which is encoded by the Locus of Enterocyte Effacement (LEE), is primarily through the master regulator Ler which is known to activate the LEE under various environmental conditions such as in response to epithelial cell contact, metabolite sensing (i.e. high acetate, low D-serine), and small intestinal pH, among others^15,16^. *C. rodentium* encodes various other factors which support gut colonization, including a Type VI Secretion System (T6SS) which it uses for interspecies bacterial competition^17^.

Previous studies examining vancomycin treatment in the context of pathogenic *E. coli* infection have focused on either prolonged antibiotic exposure prior to infection or antibiotic intervention after infection onset^18^. Consequently, little is known about the effects of brief, clinically relevant antibiotic exposure on subsequent pathogen colonization, population structure, and infection outcomes. Here, we investigate enteric infection of mice by *C. rodentium* following short-term treatment with vancomycin. We show that antibiotic pre-treatment resulted in global changes to gut colonization, with early saturation of gut sites to carrying capacity. Transcriptomic analysis indicated that increased early pathogen burden and disease severity were not driven by increased virulence, but instead reflected an altered metabolic environment that favours pathogen expansion. We investigated direct and indirect interactions between key microbiota species that bloom following the cessation of vancomycin treatment in mice. By applying a barcoding approach, we quantified how vancomycin relaxes infection bottlenecks, altering pathogen founding population size and population composition across intestinal sites. Together, these data enrich our knowledge of the relationship between enteropathogens, antibiotic use, and antibiotic-induced dysbiosis.

## Methods

### Bacterial culture conditions

All bacterial strains used in this study are summarized in Table S1. Unless otherwise stated, *C. rodentium* DBS100 ATCC 51459 was used in every experiment^19^. Prior to each assay, *C. rodentium* was streaked onto a fresh LB agar plate from frozen (−70 °C in glycerol) and grown overnight (18-20 hours) at 37 °C. A single colony was then inoculated into LB broth and grown overnight at 37 °C with agitation.

*Bacteroides* and *Akkermansia* strains were cultured in an anaerobic chamber (90% N_2_, 5% CO_2_, 5% H_2_). *B. vulgatus*, *B. ovatus*, and *B. fragilis* strains used were originally isolated from human feces^20^. *B. thetaiotaomicron* was obtained from ATCC (29148). Prior to each assay, *Bacteroides* strains were streaked onto fastidious anaerobe agar (FAA; Neogen) plates and grown overnight at 37 °C. A single colony was then inoculated into fastidious anaerobe broth (FAB; Neogen) and grown overnight at 37 °C without agitation.

Two strains of *A. muciniphila* were used in this study: *A. muciniphila* ATCC BAA-835 and human fecal isolate *A. muciniphila* CC51001 Hb generously donated by Dr. Emma Allen-Vercoe. Prior to each assay *Akkermansia* strains were streaked onto FAA supplemented with L-threonine and N-Acetylglucosamine and grown for 48 hours at 37 °C. Approximately 0.5 µL of bacteria were then inoculated into FAB modified with L-threonine and N-Acetylglucosamine and grown overnight at 37 °C without agitation.

### Animal housing and ethics

Animal experiments were performed in accordance with the Canadian Council on Animal Care and the University of British Columbia (UBC) Animal Care Committee guidelines (Animal Care Protocols A16-0216 and A20-0187). Mice were ordered from Jackson Laboratory (Bar Harbor, ME) and maintained in a specific pathogen-free facility at UBC on a 12-hour light-dark cycle, allowing for a one-week period post-arrival to acclimatize to the facility.

### Animal antibiotic treatments

One of three antibiotics was administered in the drinking water as follows: vancomycin at 100 mg/L for 2 days, metronidazole at 750 mg/L for 4 days, or streptomycin at 450 mg/L for 4 days. After antibiotic treatment all mice received standard water for a period of 24 hours to flush out any remaining antibiotic. Antibiotic concentrations were chosen as per prior animal studies^14^. While vancomycin treatment was also chosen based on a prior animal study, the chosen 100 mg/L dosage represents a low-dose exposure compared to the majority of published studies involving vancomycin treatment^18,21^.

### *C. rodentium* infections of mice

Female C57BL/6J mice were inoculated with 10^8^ CFU of wild-type (WT) or barcoded *C. rodentium* DBS100 (100 µL volume) by oral gavage. In the case of infection with the barcoded library of *C. rodentium*^22^, a 1 mL frozen aliquot of the barcoded library was subcultured for 3 hours 1:20 in LB supplemented with chloramphenicol at 30 µg/mL (+Cam30) before being spun down and washed to remove the antibiotic, and resuspended in phosphate-buffered saline (PBS). While chloramphenicol was included in the subculture media, no loss of insert stability was observed either *in vitro* or *in vivo*^22^ in the absence of antibiotic. The addition of chloramphenicol allows for consistency across studies using this barcoded library and facilitates direct comparison between studies^22^. Mice infected with the barcoded library were singly housed post-infection to prevent mixing of barcodes by coprophagy. Mice were monitored daily throughout the infection time course for weight loss and clinical symptoms. Mice were euthanized at experimental endpoint (2-7 days post-infection) by isoflurane anesthesia followed by carbon dioxide inhalation. All mice were aged 7-weeks at the time of infection and, in the case of antibiotic pre-treatment, infections were carried out one day after the removal of antibiotic treatment.

### Cecal histology

Proximal and distal cecums were fixed in 10% formalin for 24 hours before submersion in 70% ethanol. Fixed tissues were processed, paraffin-embedded, sectioned at 5 µm, and stained by Wax-it Histology Services (Vancouver, Canada). Paraffin-embedded sections were stained with hematoxylin and eosin (H&E) using standard techniques. H&E-stained sections were analyzed using a 20x lens objective on a Zeiss Axioskop 2 MOT microscope.

### Cytokine analysis

Colon samples were collected and immediately submerged in 1 mL of PBS containing protease inhibitors before homogenization in a FastPrep-24 (MP Biomedicals) at 5.5 m/s for 2 minutes. Homogenized tissues were spun down for 20 minutes at 16,000 x *g* at 4 °C and supernatants were then collected. Cytokine concentrations were determined using a mouse inflammation cytometric bead array (CBA) kit (Cat. no. 552364, BD Biosciences) and an Attune NxT flow cytometer (Thermo Fisher Scientific). Cytokine levels were normalized by tissue weight.

### Sample collection and processing for plating and barcode sequencing

Fecal samples were collected daily post-infection (p.i.). At experimental endpoint, the GI tract and spleen were collected (Figure 1C). To isolate mucosal bacterial populations, GI tissues were opened longitudinally and gently scraped to remove lumenal content. The remaining tissue was washed twice in PBS for collection, leaving only mucosal bacteria intimately associated with the epithelial tissue. All samples were collected in 1 mL PBS and homogenized in a FastPrep-24 (MP Biomedicals) at 5.5 m/s for 2 minutes. To capture as many colonizing barcodes as possible, 250 µL of the sample was plated in triplicate on MacConkey (Mac) agar +Cam30, representing 75% of the total sample. Of the remaining sample, 100 µL was diluted for plating and enumeration of colony forming units (CFU). After 18-20 hours of growth at 37 °C, bacterial colonies were counted and plates containing at least 10 colonies were scraped to collect barcoded *C. rodentium*, pooling triplicate plates, and genomic DNA was extracted (Qiagen DNeasy Blood & Tissue Kit).

**Figure 1.**
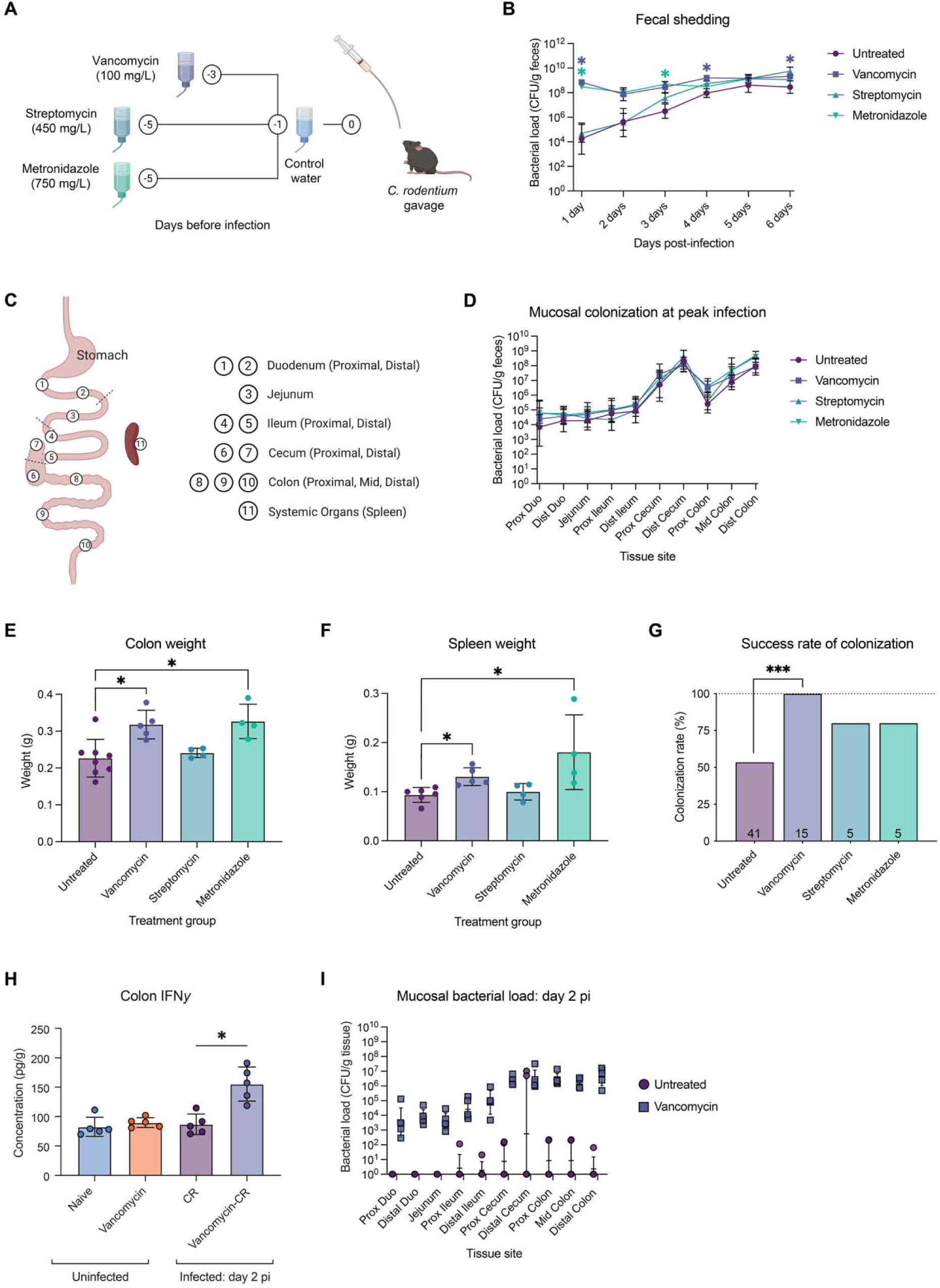
Vancomycin pre-treatment before *C. rodentium* infection dramatically alters pathogen population dynamics and associated host tissue pathology. A) Experimental setup. In brief, mice were pretreated with antibiotics in the drinking water, followed by one day on control water to flush out any remaining antibiotics, and then gavaged with *C. rodentium* at 10^8^ CFU. Antibiotics tested were: vancomycin (100 mg/L) for 2 days, streptomycin (450 mg/L) for 4 days, and metronidazole (750 mg/L) for 4 days. B) Fecal shedding per gram feces collected daily post-infection (N = 4-5). Statistics represent a Mixed-effects model with Geisser-Greenhouse correction and Dunnett’s multiple comparisons test. C) Tissue collection map of intra– and extra-intestinal sites sampled. D) Mucosal bacterial load at day 7 post-infection in untreated and antibiotic pretreated mice (N = 4-5). E) Colon tissue weight (grams) as a measure of characteristic colonic hyperplasia (N = 4-5). Statistics represent a Kruskal-Wallis test. F) Spleen weight (grams) as an indicator of host inflammatory state (N = 4-5). Statistics represent a Kruskal-Wallis test. G) Success rate of *C. rodentium* colonization in untreated mice compared to those pre-treated with either vancomycin, metronidazole, or streptomycin, as measured by the number of singly-housed mice with robust colonization as of day 4 post-infection. Statistics represent a Fisher’s exact test. H) IFNγ cytokine levels (pg/g) in uninfected mice, vancomycin pretreated mice, and at day 2 pi with or without antibiotic pretreatment (N = 5). CR, *C. rodentium*. Statistics represent a Kruskal-Wallis test with Dunn’s multiple comparisons test. I) Mucosal bacterial load at day 2 pi in untreated-infected and vancomycin-infected mice (N = 5).

### Cecal content collection and processing of supernatants

Cecal contents were collected from either untreated mice or mice pre-treated with vancomycin as described above. In the case of vancomycin-treated samples, contents were collected one day following antibiotic cessation. Gut contents were homogenized in 1:2 w/v reduced PBS using a Mixer Mill MM 400 (Retsch) for 5 minutes at an oscillation frequency of 25. Samples were then spun at 7500 rpm for 10 minutes for supernatant collection. Supernatants were pooled from 4-5 mice to control for differences between individuals and ensure consistent results. Separate supernatant pools were used for each biological replicate presented in this study.

### RNA isolation and sequencing

Wild-type *C. rodentium* DBS100 (CmR) was cultured to mid-log phase (3.5 hours) in 1:1 DMEM: Cecal supernatant from naïve or vancomycin-treated mice in a 96-well plate at 37 °C with agitation. *C. rodentium* cultured in DMEM or DMEM + 30 µg/mL of chloramphenicol were included as controls. Bacterial cells were collected, treated with RNAprotect Bacterial Reagent (Qiagen), and stored at –70 °C until extraction. RNA was isolated using the RNeasy Mini Kit (Qiagen), and genomic DNA was digested on column using an RNase-Free DNase Set (Qiagen). RNA quality was assessed using a 2100 Bioanalyzer (Agilent Technologies, USA) to ensure an RNA Integrity Number (RIN) higher than 6. A RIN score of 6 is considered high quality for *C. rodentium*^23^. Due to the potential for cecal supernatants to contain exogenous RNA, uninoculated blank samples were included, extracted, and assessed via Bioanalyzer. We did not observe peaks corresponding to bacterial RNA in these blank samples, indicating that the RNA obtained and sequenced originated only from inoculated *C. rodentium*.

Library preparation, rRNA depletion, and sequencing were performed by the Genome Sciences Centre at the BC Cancer Research Centre. In brief, rRNA depletion was performed using the bacterial NEBNext rRNA Depletion Kit (Cat. no. E7850, New England BioLabs). Sequencing libraries were created by end repair and phosphorylation, followed by 3′ A-addition and adapter ligation using a custom reagent formulation (Cat. no. E6000B-10, New England BioLabs). Libraries were pooled in equal molar amounts and sequenced on an Illumina HiSeqX platform (Illumina) with paired-end (PE) reads.

### RNA-sequencing analysis and pathway enrichment

cDNA sequencing reads were aligned to the *C. rodentium* ICC168 reference genome (NC_013716, NC_013717, NC_013718 and NC_013719)^24,25^ and counts were generated by using STAR 2.7.10a^26^ (v2.7.10a). Quality of the sequencing reads was assessed using FastQC^27^ (v0.11.9), alignment was assessed using RSeQC^28^ (v4.0.0), and quality control reports were assembled using MultiQC^29^ (v1.12). At least 10 million PE reads were obtained per sample. Reads were filtered to include transcripts present in at least 3 samples with a count of 10 or higher for analysis. DESeq2^30^ was used to identify differentially expressed genes (DEGs) using Independent Hypothesis Weighting (IHW)^31^ for *p*-value adjustment and the ‘ashr’ package for LFC shrinkage^32^. Differential regulation was defined as genes with an adjusted *p*-value of < 0.05 and a log2 fold change equal to or greater than ± 1.5 when comparing *C. rodentium* in untreated vs vancomycin-treated cecal supernatants.

Pathway enrichment analysis was performed by using Cytoscape v3.9.1^33^ using a separate pre-ranked list of up– and downregulated genes against the background universe of all detected genes (after pre-filtering). Genes were arranged according to log2FC. Pathways were determined according to Gene Ontology (GO) terms^34^ in the GO Biological Processes database, with redundant GO terms removed using a redundancy cut-off of 0.5. Pathways with an FDR < 0.05 were considered significantly enriched.

### HT29-MTX cell culture assays

HT29-MTX human colonic cell line (European Collection of Authenticated Cell Cultures; Cat. No. 12040401) was grown in DMEM (Hyclone) supplemented with 1% v/v non-essential amino acids (Gibco, Cat. No. 11140-050), 1% v/v Glutamax (Gibco, Cat. No. 35050061), 10% v/v heat-inactivated fetal bovine serum (FBS) and 100 U/mL PenStrep (Gibco, Cat. No. 15140122) at 37 °C and 5% CO_2_. Cells were passaged depending on their confluency every 2-3 days and seeded into plates at a density of 1.25 x 10^5^ cells/mL after at least 4 routine passages. Cells were allowed to differentiate for 21 days as previously described, to ensure robust mucus production, with media changed every 2 days^35,36^.

After differentiation, cell culture media was removed and replaced with fresh media equilibrated at 37 °C and 5% CO_2_ and inoculated with *C. rodentium* at a multiplicity of infection (MOI) of 100. Inoculated plates were spun at 1,000 rpm for 5 minutes to synchronize the infection before incubation at 37 °C and 5% CO_2._ After a 4-6 hour infection, culture supernatants were collected by pipetting and diluted for plating on selective media. Remaining cells were washed 5 times to remove unattached bacteria before detaching from the plating using 0.1% Triton X-100 in PBS. Detached cells were collected for dilution and plating on selective media. Resultant colonies were counted for the enumeration of CFU.

In the case of *A. muciniphila* and *Bacteroides* co-inoculation, commensal bacteria were inoculated for a period of 1 hour before the addition of *C. rodentium* to allow for pre-processing of the mucus layer. *A. muciniphila* and *Bacteroides* strains were inoculated 1:10 from an overnight culture at an OD of 0.5-0.8.

### 16S rRNA microbiota sequencing

Fecal pellets were collected from mice and stored at –70 °C until extraction. DNA extraction was performed using a Qiagen PowerFecal kit. The V4 variable region of the 16S rRNA gene was amplified using indexed 515F 806R primers (Table S2). Indexed PCR products were pooled and sequenced on a MiSeq platform (Illumina) using V2 technology. Raw reads were quality-filtered and processed in QIIME2 using DADA2. Taxonomy was assigned using a Naïve Bayes classifier trained on the SILVA 138 database for the 515/806 region, with a 99% OTU cutoff. Further filtering was performed in R using phyloseq (v1.38.0)^37^ to remove singletons and rarefy samples to 4800 reads. Alpha and beta diversity analyses were performed using phyloseq and plotted with ggplot2.

### Reporter strain construction and assays

To generate the *C. rodentium* reporter strains, the promoters upstream of *ler* and *tir* were identified using convolutional neural network (<u>CNN</u>) and neural network promoter prediction (<u>NNPP</u>) servers^38,39^. Gene sequences were synthesized by Genewiz. The fragments containing regulatory regions were inserted into the PmeI and SnaBI sites in pGEN-luxCDABE (Addgene; #44918). *C. rodentium* was transformed using electroporation, and bacteria harbouring the plasmid were selected using carbenicillin (100 µg/mL).

Virulence reporter assays in response to *Bacteroides* were performed by inoculating either the Ler or Tir reporter strains 1:100 into FAB containing either of *B. thetaiotaomicron*, *B. ovatus*, *B. vulgatus*, or *B. fragilis* 1:10 from overnight culture, or supernatants of overnight culture from the same four strains. Assays were performed in a black, clear-bottom 96-well plate incubated at 37 °C for 6 hours under anaerobic conditions without agitation. After 6 hours, luminescence was read on a Synergy HI plate reader (Biotek) under aerobic conditions. Cultures were then diluted and plated on MacConkey agar for CFU. Luminescence readouts were normalized to *C. rodentium* CFU. *C. rodentium* cultured in DMEM under the same conditions was included as a control, as DMEM is known to activate expression of virulence genes.

Virulence reporter assays in response to *A. muciniphila* were performed as described above, but with inoculation into modified FAB (See ‘Bacterial culture conditions’) to better support the growth of *A. muciniphila*. Assays were only performed using the *A. muciniphila* ATCC BAA-835 strain.

### Metabolite extractions and data analysis

Cecal contents were collected from naïve and vancomycin-treated mice for untargeted metabolomics analysis. 10 mg of the sample was weighed for extraction. 1000 µL 1:4 v/v LC-MS grade MeOH:water was added before disruption by water bath sonication. Sample supernatant was collected by centrifugation, air-dried in a speed vac and stored at –70 °C until use. Dry samples were reconstituted according to sample weights at a concentration of 10 mg/100 µL. LC-MS was conducted as a commercial service by Dr. Tao Huan’s group (University of British Columbia) using a UHR-QqTOF (ultrahigh resolution Qq-time-of-flight) mass spectrometer and sample concentrations were normalized by weight. Identified metabolite concentrations were analyzed using Metaboanalyst 5.0 (www.metaboanalyst.ca). Features with a constant or single value across samples were removed. Data were further filtered to remove features with a high relative standard deviation (low repeatability) of > 20 %. Samples were normalized by median before log transformation and scaling based on the dispersion of each feature. Metabolites were annotated by mass/charge (m/z) ratio using The Human Metabolome Database (HMDB) with a molecular weight tolerance of 5 ppm.

### Multi-omics integration of transcriptomics and metabolomics data

Individual weighted gene correlation networks were constructed using WGCNA (v1.73)^40,41^. The relationships between dataset modules, and between modules and treatment groups, were calculated by Pearson correlation. Triangular transcript-metabolite-trait relationships were mapped using igraph (v2.1.4)^42^ and further prepared for visualization using Cytoscape v3.9.1^33^.

### Bacterial growth curves

*C. rodentium* growth was assessed by inoculating 1:200 from an overnight culture into LB media. OD 600 nm measurements were taken at 10-minute intervals using a Synergy H1 plate reader (Biotek) at 37 °C with agitation.

Daidzein (Thermo Scientific; Cat. no. AC328230250) and equol (Thermo Scientific; CAS Number 94105-90-5) were supplemented into LB media by adding 150 µM of either metabolite, with concentrations chosen to be physiologically relevant according to ^43^. Growth was assessed as described above.

Bile acid supplementation was carried out as follows: Separate 10X stock solutions of cholic acid, deoxycholic acid, and lithocholic acid were prepared in methanol. Bile acids were supplemented into LB culture media to a final concentration of 1 mg/mL, representing the estimated levels in the cecum and colon. LB supplemented with methanol was included as a vehicle control at corresponding volumes. Growth was assessed by dilution and plating on LB agar after 20 hours in liquid culture at 37 °C with agitation.

### Microbial cross-feeding

Co-culture assays were performed as follows: *C. rodentium* and *A. muciniphila* strains were cultured as described above before inoculation into mucin-rich media^44^. *A. muciniphila* strains were inoculated 1:20 from an overnight liquid culture. *C. rodentium* was inoculated 1:200 from an overnight liquid culture. Cultures were grown for 24 hours before dilution and plating onto MacConkey agar (Difco) which is selective for *Enterobacteriaceae*.

To test for cross-feeding of mucin cleavage products, *C. rodentium* was grown in spent media from *A. muciniphila* culture. To prepare these spent media, *A. muciniphila* was inoculated 1:20 from an overnight culture as before, into media containing mucin as the sole carbon source (mucin-only media): 10X M9 salts^45^ supplemented with 1% type II porcine stomach mucin (Sigma-Aldrich). Cultures were grown for 24 hours before spinning at 3200 rpm for 15 minutes to pellet the bacteria and collect the resultant supernatants for *C. rodentium* inoculated at 1:200 for 24 hours before dilution and plating. Samples were plated both on MacConkey agar for enumeration of *C. rodentium* CFU and on modified FAA agar to confirm the absence of *A. muciniphila*. Supernatant from mucin-only media not inoculated with *A. muciniphila* was included as a control. A starvation control of mucin-only media inoculated with water was included to represent nutrient depletion by *A. muciniphila*.

### Sequencing library preparation for barcode sequencing

Preparation of the library for amplicon sequencing involved the addition of index tags by triplicate PCR to reduce PCR bias as described^22^. Cycling conditions were as follows: initial 30 s denaturation at 98 °C followed by 30 cycles of 10 s at 98 °C, 10 s at 52 °C, and 28 s at 72 °C with a 5 min final extension at 72 °C. Triplicate PCR reactions were pooled and purified using the Qiagen MinElute PCR Purification Kit before DNA quantification using a Quant-iT PicoGreen dsDNA Assay Kit (Invitrogen). Equal concentrations of each purified sample were then pooled and sequenced.

### Amplicon sequencing of barcoded *C. rodentium* and data analysis

Sequencing was performed on a MiSeq (Illumina) using a 50-cycle V2 MiSeq reagent kit with 40 % PhiX and a custom read 1 sequencing primer^22^. All sequencing was done at the UBC Biomedical Research Centre Sequencing Core. Demultiplexed single-end reads were processed in QIIME2, trimming conserved regions and filtering for barcodes of exactly 30 bps in length. Denoising was performed using DADA2 to distinguish sequencing errors from true biological variability, and thereby infer exact amplicon sequence variants (ASVs). Data were then filtered to include barcodes present in a minimum of 5 reads in at least one endpoint sample, and present in the inoculum in at least 2 reads. The library has been previously defined as including an average of 2,062 unique barcodes^22^. Previously described formulas were used to calculate the estimated population size (Nb), allele frequency, genetic relatedness, and directionality index^46–48^. Alleles were substituted for individual barcodes as described (5), and generation time (g) was set to 1 for all calculations. Calculated Nb values were corrected using an *in vitro* calibration curve (IVCC) previously generated and reported by ^22^. Corrected values were plotted as Nb’. In directionality index calculations, the number of unique inoculated barcodes was used to determine population size (n). Untreated *C. rodentium* population dynamics were calculated using the published NCBI Sequence Read Archive (SRA) dataset BioProject ID PRJNA810564^22^, as the experiments shared control animal subsets.

### Statistical analysis

Statistical analysis was performed in GraphPad Prism (www.graphpad.com). All statistical tests used are specified in their respective figure legends. Unless otherwise stated, analysis of non-normally distributed data was performed using a Mann-Whitney U test to compare two groups and Friedman test for more than two groups with Dunn’s multiple comparisons test. Unless otherwise stated, analysis of normally distributed data was performed using two-way ANOVA with Dunnett’s multiple comparisons test, or a Mixed-effects analysis with Holm-Šidák’s multiple comparisons test. Aggregate results represent the mean +/-SD, and statistical significance is represented by \**p* < 0.05, \*\**p* < 0.01, \*\*\**p* < 0.001 and \*\*\*\**p* < 0.0001. Spearman correlation analysis between variables was performed in GraphPad Prism (*p*– and rho-values representing Spearman correlation values; lines representing linear regression +/− 95% confidence intervals). Non-metric Multi-Dimensional Scaling (NMDS) plots were created by calculating the Bray-Curtis distance matrix using phyloseq (1.38.0)^37^.

### Data visualization

Data visualization was performed in R (4.1.0) using the packages phyloseq (v1.38.0)^37^, tidyverse (v1.3.1)^50^, ggplot2 (v2.3.3.5)^51^, ggrepel (v0.9.1)^52^, pheatmap (v1.0.12)^53^, and ggsci (v2.9)^54^.

## Results

### Vancomycin pre-treatment dramatically alters *C. rodentium* colonization dynamics and exacerbates colitis pathology

Given the importance of colonization resistance to infection susceptibility, we first profiled the colonization dynamics and disease course of *C. rodentium* in the context of antibiotic-induced gut dysbiosis. We tested the effects of pre-treatment with either vancomycin, streptomycin, or metronidazole on temporal and regional gastrointestinal colonization by *C. rodentium*. Streptomycin and metronidazole treatments were chosen to target and reduce colonization by a broad spectrum of microbes, and have been previously investigated in the context of pre-treatment followed by *C. rodentium* infection, with metronidazole increasing the severity of subsequent colitis symptoms^14^. Vancomycin was further chosen to target Gram-positive bacteria.

Following antibiotic administration in the drinking water, all mice were returned to regular drinking water for a period of one day before *C. rodentium* inoculation (Figure 1A). We first assessed fecal shedding, typically characterized by initial low-level shedding (10^3^-10^4^ CFU per gram feces at 1 day post-infection [p.i.]), which steadily increases towards peak infection^22^. Following antibiotic treatment, we noted higher fecal shedding in vancomycin– and metronidazole-treated mice with up to a 6-log increase in fecal burden at day 1 p.i. compared to untreated and streptomycin-treated mice, indicative of altered early infection events (Figure 1B). However, tissue-resident pathogen burden (Figure 1C) at peak infection on day 7 was the same across all groups (Figure 1D), though vancomycin did result in a trend towards higher colonization of the proximal colon.

Despite saturation of *C. rodentium* burden within the gut at peak infection, differences in early colonization dynamics may still influence overall disease outcome^22^. Indeed, colon weight at peak infection, an indicator of colonic hyperplasia, was increased in response to both vancomycin and metronidazole, as was spleen weight, indicating increased systemic inflammation (Figure 1E-F). At the population level, the overall success rate of colonization, determined by the number of singly housed mice exhibiting *C. rodentium* colonization 7 days post-inoculation, increased from 53% in non-antibiotic-treated singly housed mice to 80% in streptomycin and metronidazole conditions, and 100% in vancomycin pre-treated mice (Figure 1G). Vancomycin also impacted systemic colonization, with both a higher systemic CFU burden and a higher rate of developing systemic infection in gut-colonized mice, increasing from 81% in untreated mice to 100% in the vancomycin pre-treatment group (Figure S1A-B). Together, these data suggest that infection dynamics within the vancomycin-induced gut environment are significantly altered with consequences to host disease outcome.

To date, the only investigation of vancomycin treatment and *C. rodentium* infection has been done in the context of antibiotic intervention following established infection^18^ rather than antibiotic pre-treatment as in this current study. Considering the evidence for altered infection dynamics we sacrificed mice at day 2 p.i. to evaluate changes to inflammation and colonization during early infection within the vancomycin-induced environment. Despite day 2 being early in infection, characterized by variable colonization under typical infection conditions^22^, we found a significant increase in levels of inflammatory cytokine IFNγ by cytometric bead array (CBA) in colonic tissues of vancomycin pre-treated infected mice, while observing no elevated IFNγ in response to either vancomycin or *C. rodentium* alone (Figure 1H). These results corresponded to a high mucosal pathogen burden across all gut regions at levels comparable to peak infection (Figures 1I and 1D). As expected, naïve-infected mice were only robustly colonized within the distal cecum, which houses the cecal patch^55^. At day 7 p.i. we further observed tissue pathological changes. Infection caused by *C. rodentium* results in colitis characterized by hyperproliferation of cells within the colonic crypt, leading to thickening of the intestinal epithelium. At peak infection gross pathological changes were noted in the cecum and proximal colon of infected mice pre-treated with vancomycin compared to untreated mice, suggesting changes to this hyperproliferation phenotype. Histological analysis revealed increased proximal cecum thickness as a result of combined vancomycin pre-treatment and *C. rodentium* infection, but not in either condition alone (Figure S1C).

### *C. rodentium* downregulates expression of key metabolism and virulence genes in response to the vancomycin-induced cecal environment

Given that colonization, inflammation and tissue pathology are significantly altered by vancomycin pre-treatment, we next sought to determine whether the behaviour and virulence of *C. rodentium* is influenced by the vancomycin-induced gut environment, allowing for increased early attachment across gut tissues. This was done by culturing *C. rodentium* in gut supernatants collected from either untreated or vancomycin-treated mice, before the assessment of gene expression via RNA-seq (Figure 2A). We focused on the supernatant from processed cecal content, as the cecum represents the site of initial *C. rodentium* colonization^55^. To investigate changes to bacterial gene expression, we performed transcriptomic analysis on *C. rodentium* grown to mid-log phase in 1:1 cecal supernatant and DMEM base media. Principal component analysis revealed differences in the clustering of *C. rodentium* transcriptomic profiles in response to untreated and vancomycin pre-treated cecal supernatants (Figure S2A), indicating altered gene expression profiles. We identified 104 differentially expressed genes (DEGs) between the two groups (Supplementary dataset 1). DEGs were identified using a minimum fold change cut-off of 1.5, with the majority being downregulated in response to vancomycin-treated cecal supernatants (83 DEGs; Figure 2B). These included operons involved in cobalamin (vitamin B12) biosynthesis (*cbiBDGM*)^56^, and structural and accessory genes of the citrate lyase complex (*citCDEFGTX*), which have been shown to regulate acid tolerance in response to fermentation products^57^. Comparatively few upregulated DEGs were identified (21 DEGs), most of which were involved in ethanolamine utilization (*eutDEGJMNPTQ*)^58^. Enrichment analysis of Gene Ontology (GO) terms showed a significant downregulation of metabolic pathways involved in cobalamin biosynthesis, small molecule metabolism, aldonic acid catabolism, methylation and nitrogen metabolism (Figure 2C). Significantly upregulated pathways included formate metabolic processes, putrescine transport, and cellular processes. The presence of the alcohol catabolism pathway as both up– and downregulated is likely reflective of the upregulation of ethanolamine utilization genes and downregulation of propanediol utilization genes (*pduABJKLNPTUW*)^59^. Together, these data indicate major shifts in pathogen metabolism in response to the vancomycin-induced cecal environment.

**Figure 2.**
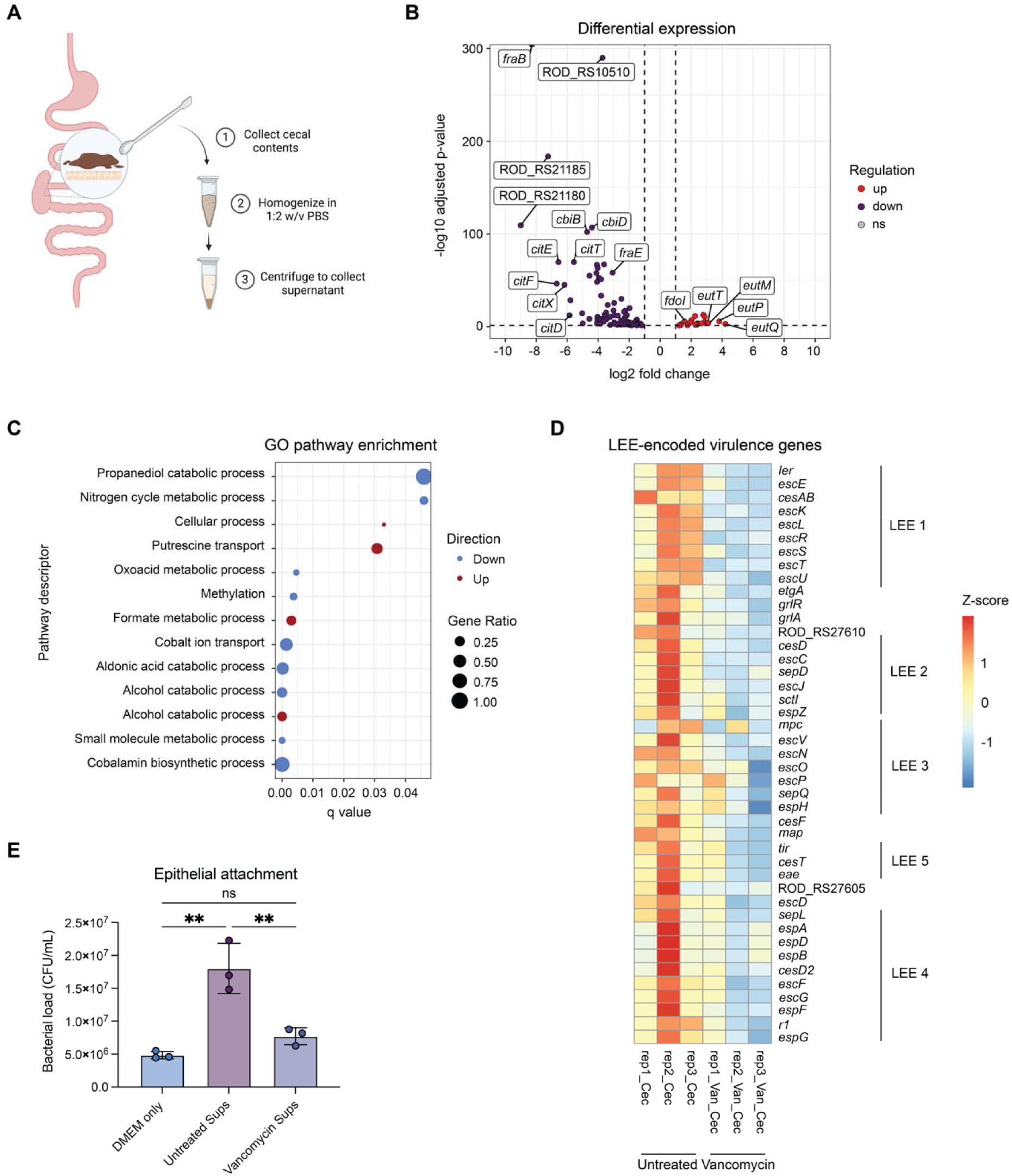
*C. rodentium* alters its metabolism in response to the vancomycin-induced gut environment, downregulating virulence. A) Outline of cecal supernatant collection and processing for use in *in vitro* assays. B) Volcano plot of differentially expressed genes (DEGs) with a fold change cut-off of 1.5 and *p*-value <0.05. C) GO pathway enrichment analysis of DEGs compared to a universe of 4928 genes. D) Heatmap showing Z-score expression of genes encoded in the Locus of Enterocyte Effacement (LEE) associated with the type III secretion system. Vertical lines indicate the operon (LEE 1-5). E) Attachment of *C. rodentium* to an epithelial cell line (HT29-MTX) after pre-induction to mid-log phase in DMEM base media or DMEM base media supplemented 1:1 with either untreated or vancomycin-treated cecal supernatants. N = 3 biological replicates (average of 3 technical replicates each), with each biological replicate performed in a unique pool of supernatants from 4-5 mice each. Statistics represent an Ordinary one-way ANOVA with Tukey’s multiple comparisons test.

Metabolism is directly linked to the expression of virulence in enteric pathogens^60^. Ethanolamine in particular, known to be present within the gastrointestinal tract, has been shown to both act as a nitrogen source for related pathogen *E. coli* O157:H7 and to stimulate an upregulation of virulence^58,61–63^. In *C. rodentium*, the transcription factor EutR senses ethanolamine in the environment leading to EutR-dependent activation of the LEE^64^. Therefore, we next wanted to determine whether the increased disease severity and colonization of diverse founders observed during infection of vancomycin-treated mice was due to differential regulation of key virulence machinery. We analyzed the expression of key virulence genes associated with the type III secretion system^65,66^ and type VI secretion systems (T6SS; CTS1 and CTS2)^17^. While we expected an upregulation of LEE-encoded genes, we observed a trend towards decreased T3SS-related gene expression in response to cecal supernatants from vancomycin-treated mice as compared with controls (Figure 2D). We further expected to see changes in the regulation of either of *C. rodentium*’s two T6SSs, which are important tools during competition with other microbes^17^. However, we did not observe any differences in either system as a result of antibiotic exposure (Figure S2B-C).

To determine how these changes in transcriptomic regulation impact virulence behaviours, we again pre-induced *C. rodentium* in cecal supernatant from either untreated or vancomycin-treated mice. This time however, *C. rodentium* cultures were washed and resuspended in cell culture media before being used to infect HT29-MTX cells, a mucus-producing colonic cell line, to assess epithelial cell attachment. After a 4-hour infection period, cells were washed to remove non-adherent bacteria before plating of the attached population. We found that attachment of *C. rodentium* pre-induced in supernatants from vancomycin-treated mice was significantly reduced as compared to pre-induction in supernatants from untreated mice (Figure 2E), consistent with the RNA sequencing results. It is clear that the relationship of *C. rodentium* with the vancomycin-induced gut environment is more nuanced than just impacting pathogen virulence to lead to more severe disease.

### Spatio-temporal interactions between *C. rodentium* and the vancomycin-associated microbiota contribute to intestinal pathogen colonization

Re-colonization of vancomycin-treated mice with an intact specific pathogen-free (SPF) microbiota via FMT was sufficient to restore pathogen colonization to naïve-infected levels (Figure 3A). Therefore, the widespread metabolic changes observed in our pathogen transcriptomic analysis likely reflect a change in microbiota-host or microbiota-pathogen interactions following antibiotic treatment, thereby allowing for robust pathogen expansion. To investigate specific pathogen-commensal relationships, we first needed to characterize the vancomycin-associated microbiota encountered by *C. rodentium* upon infection. As expected, 16S rRNA sequencing on fecal samples from uninfected animals before, during, and after antibiotic treatment demonstrated a significant decrease in species richness post-antibiotic treatment (Figure S3A), and principal coordinate analysis (PCoA) demonstrated a clear separation in Bray-Curtis distance between pre-treatment groups (Figure S3B). Vancomycin treatment resulted in a significant loss of *Lactobacillaceae*, *Muribaculaceae, Rikenellaceae, Lachnospiraceae,* and *Erysipelotrichaceae* (Figure 3B). The vancomycin-associated microbiota was instead dominated by two bacterial genera: *Bacteroides* and *Akkermansia* (Figure 3C). *Akkermansia* bloomed during antibiotic treatment, increasing from an average relative abundance of 3.1% before antibiotic treatment to 48.7% at the time of *C. rodentium* gavage. In contrast, *Bacteroides* bloomed following the cessation of antibiotic treatment, increasing from an average relative abundance of 5% before and during antibiotic treatment to 45.7% at the time of *C. rodentium* gavage (Figure 3C). The next most abundant genus was *Parasutterella* at only 3.45% abundance. This demonstrates that *C. rodentium* interacts with a bacterial community with limited diversity during early colonization of the vancomycin-perturbed gut.

**Figure 3.**
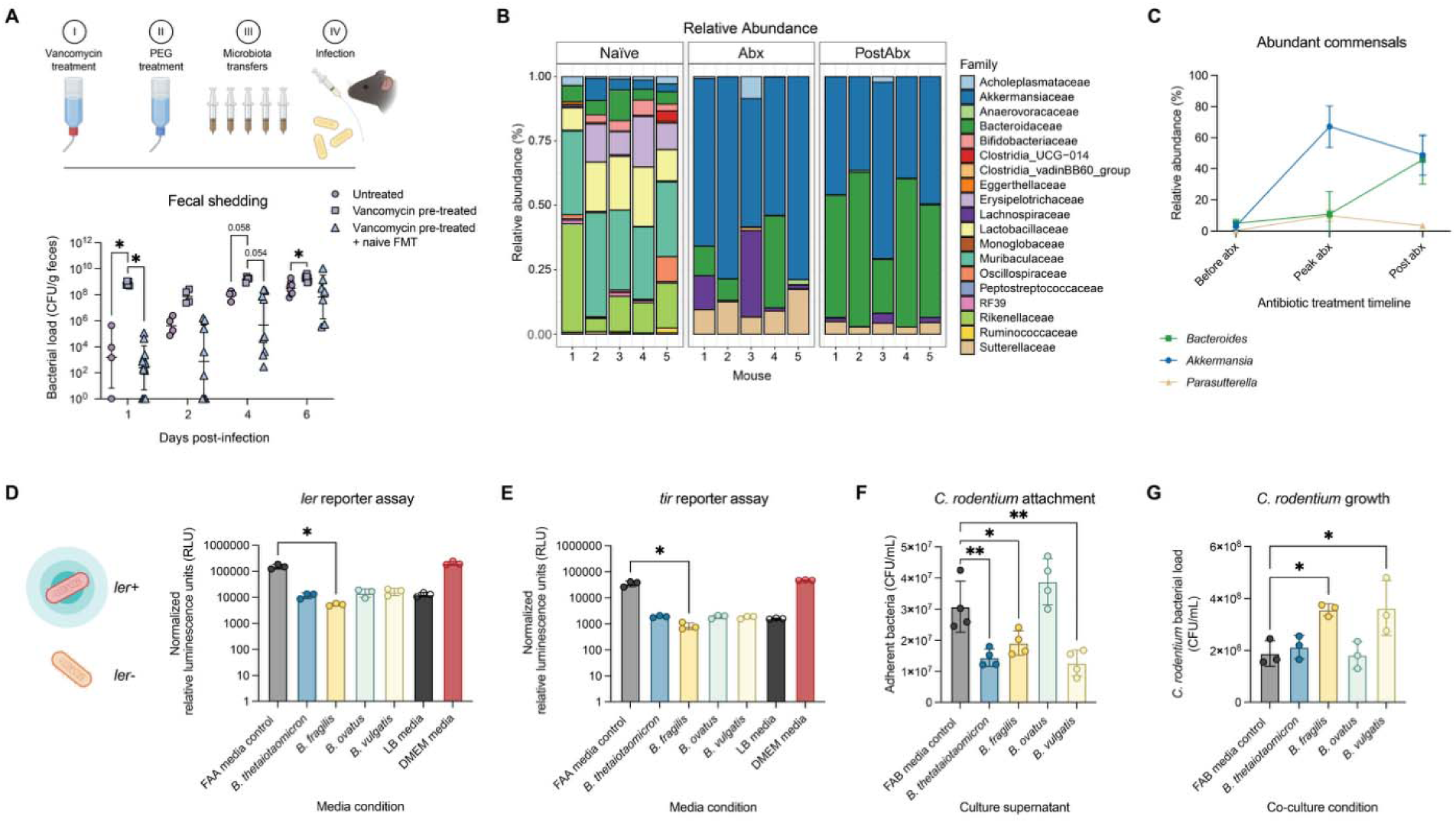
The vancomycin-associated microbiota impacts growth and epithelial cell attachment of *C. rodentium*. A) Outline of fecal microbiota transfer (FMT) experiments, detailing the transfer of a naïve-associated microbiota to vancomycin-treated mice over the course of six microbial transfers followed by *C. rodentium* gavage. Here we show shedding of *C. rodentium* in the feces of infected mice following no-treatment, vancomycin-treatment, or vancomycin-treatment with FMT from untreated mice (N = 4-12). Statistics represent a Mixed-effects model with Geisser-Greenhouse correction and Tukey’s multiple comparisons test. B) Family-level relative abundance (%) of the murine microbiota sampled before vancomycin treatment (Naïve), after 2 days on vancomycin (Abx), and one day following cessation of vancomycin treatment (PostAbx) (N = 5). Temporal data is matched by mouse. C) Top 3 most abundant commensal microbes sampled before vancomycin treatment (Before abx), after 2 days on vancomycin (Peak abx), and one day following cessation of vancomycin treatment (Post abx) (N = 5). D) *C. rodentium ler* gene expression as measured by relative luminescence using a *ler* reporter strain. E) *C. rodentium tir* gene expression as measured by relative luminescence using a *tir* reporter strain. Data normalized to *C. rodentium* CFU. F) Adherent *C. rodentium* after 6-hour infection of HT29-MTX colonic epithelial cells in response to *Bacteroides* culture supernatants. G) *C. rodentium* growth after 24 hours in co-culture with *Bacteroides* strains.

Both *B. thetaiotaomicron* and *A. muciniphila* have been associated with increased severity of *C. rodentium* infection in naïve SPF mice^67,68^. To determine their impacts on virulence regulation, we first evaluated expression of *C. rodentium* virulence genes in response to *Bacteroides* and *Akkermansia*. To do this we used *ler* and *tir* reporter strains of *C. rodentium*, which include a luciferase cassette downstream of the native promoter for each gene, causing the bacterium to luminesce proportionally to gene expression. Using these reporter strains, we identified a decrease in expression of both *ler* and *tir* in response to supernatants from four commensal *Bacteroides spp.* (*B. thetaiotaomicron*, *B. fragilis, B. vulgatus,* and *B. ovatus*) as compared to the media control (Figure 3D-E), here presented as luminescence normalized to *C. rodentium* CFU. In comparison, supernatants from two isolates of *Akkermansia muciniphila* did not alter expression of either *ler* or *tir* (Figure S3C-D). This suggests that neither *Bacteroides* nor *Akkermansia* directly stimulate the expression of attachment-related genes, and that *Bacteroides* may inhibit expression of attachment-related genes.

To determine how the presence of metabolites from *Bacteroides* could impact pathogen attachment to epithelial cells, we infected the mucus-producing epithelial cell line HT29-MTX with *C. rodentium* with or without the addition of supernatants from *Bacteroides* cultures. We found that supernatants from *B. thetaiotaomicron*, *B. fragilis*, and *B. vulgatus* significantly decreased pathogen adherence to HT29-MTX cells after a 6-hour infection (Figure 3F). We also found that planktonic *C. rodentium* CFU were increased in the same wells (Figure S3E). *Bacteroides* are known to be highly metabolically active. For example, *B. thetaiotaomicron* is known to increase the concentration of freely available sugars which can promote pathogen growth^69^. Therefore, an overabundance of *Bacteroides* may similarly allow for population expansion by *C. rodentium* during colonization. We investigated the growth of *C. rodentium* either alone or in co-culture with *Bacteroides* and found that co-culture with *B. fragilis* and *B. vulgatus* both significantly increased *C. rodentium* growth after 24 hours (Figure 3G).

### Differentially abundant metabolites in the vancomycin-induced gut environment impact pathogen expansion

The disparate relationship between increased early mucosal colonization and decreased virulence gene expression and epithelial cell attachment prompted us to further investigate the relationship between key metabolites, commensal microbes, and *C. rodentium*. By performing untargeted metabolomics on cecal contents of untreated and vancomycin-treated mice, we found distinct clustering of metabolic profiles by principal component analysis (PCA), indicating a substantial change to cecal metabolomic composition (Figure S4A). Indeed, we found that 1728 metabolites were differentially abundant in cecal contents following vancomycin treatment when compared to untreated controls. Of these metabolites, 857 increased in abundance and 847 metabolites decreased in abundance in the vancomycin-induced cecal environment (Figure 4A; Supplementary dataset 2).

**Figure 4.**
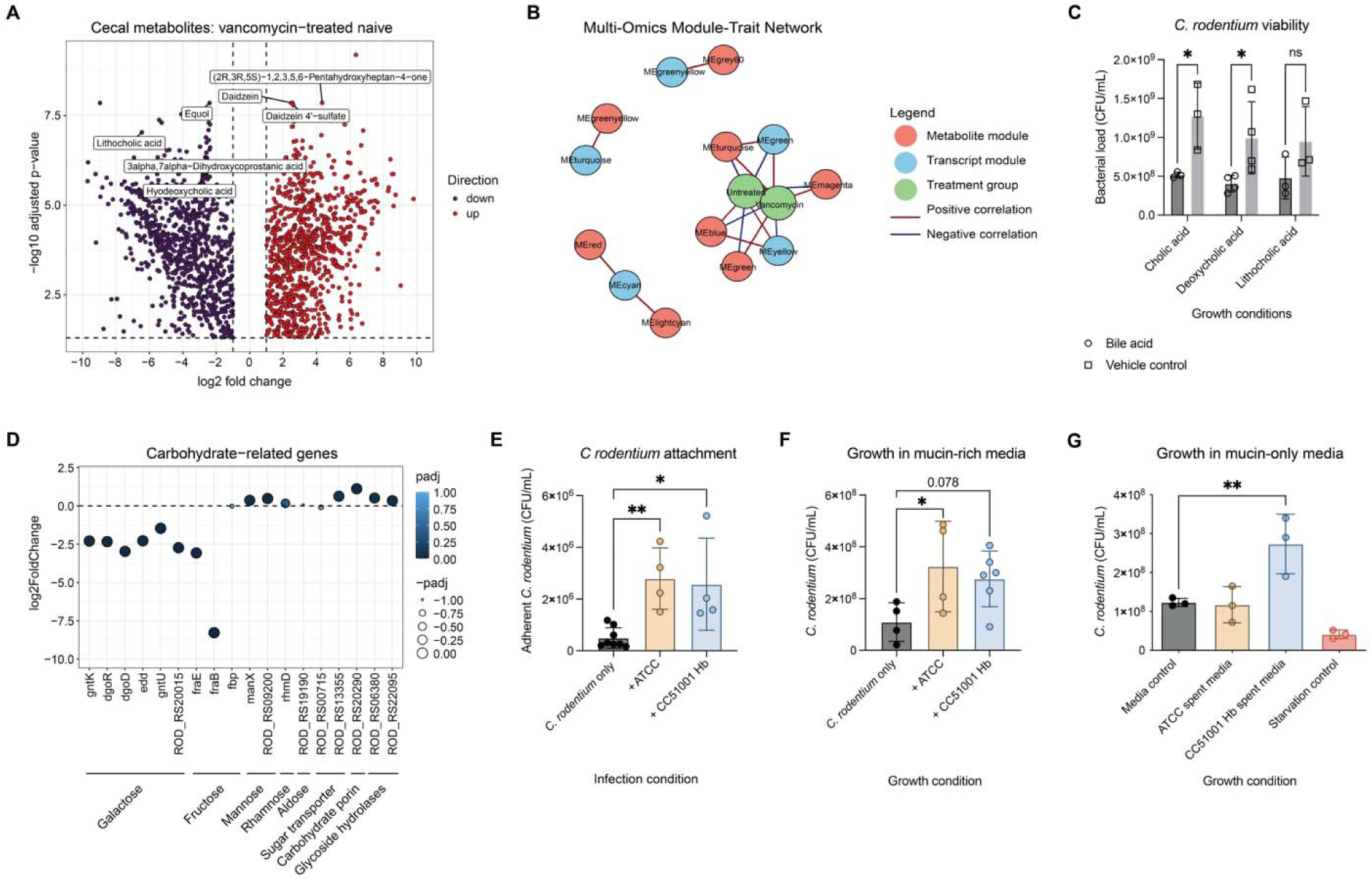
Metabolomic shifts in the vancomycin-treated gut support pathogen expansion. A) Volcano plot of differentially abundant metabolites with a log2 fold change cut-off of 2 and *p*-value <0.05. B) Network analysis relating transcript and metabolite correlation networks to treatment conditions. C) Growth of *C. rodentium* in LB media supplemented with 1mg/mL of either cholic acid, deoxycholic acid, lithocholic acid, or methanol vehicle control for 24 hours (N = 3 biological replicates). D) Log2 fold change of *C. rodentium* carbohydrate-related genes following growth in cecal supernatant from vancomycin-treated mice compared to untreated mice. E) Adherent *C.* rodentium after 6-hour infection of HT29-MTX epithelial cells in co-culture with *A. muciniphila*. G) Growth of *C. rodentium* after 24 hours of anaerobic growth in co-culture with *A. muciniphila* strains in mucin-rich media. H) Growth of *C. rodentium* after 24 hours of anaerobic growth in spent media from *A. muciniphila* strains grown in mucin as a sole carbon source. E-H) Statistics represent a one-way ANOVA with Holm-Šidák’s multiple comparisons test. Each data point represents a biological replicate (the average of three technical replicates).

To further link metabolic signatures to *C. rodentium* gene expression, we performed weighted gene correlation network analysis (WGCNA) of gene expression and untargeted metabolomic data, allowing us to identify triangular transcript-metabolite-trait relationships (Figure 4B; Figure S4B). We were able to identify two such triangular relationships: (1) a positive correlation between transcript module MEgreen (270 genes), metabolite module MEturquoise (1189 metabolites) and vancomycin treatment; and (2) a positive correlation between transcript module MEyellow (293 genes) and metabolite module MEblue (463 metabolites) which were both negatively correlated to the vancomycin treatment condition (Supplementary dataset 3).

Many metabolites that decreased in abundance included those processed by commensal gut microbes, such as select bile acids (lithocholic acid, hyodeoxycholic acid, 3alpha,7alpha-dihydroxycoprostanic acid, and cholic acid) (Figure 4A; Figure S4C). The metabolite module MEblue, which negatively correlated with vancomycin treatment, was enriched for bile acids, including deoxycholic acid and taurocholic acid. Considering that *C. rodentium* is not able to colonize the gallbladder, which stores bile acids^22^, we hypothesized that the loss of specific metabolites following vancomycin treatment could promote *C. rodentium* colonization. We therefore investigated whether bile acids are capable of inhibiting *C. rodentium* growth. We cultured *C. rodentium* in LB supplemented with three bile acids which decreased in abundance following vancomycin treatment: primary bile acid cholic acid and secondary bile acids deoxycholic acid and lithocholic acid. Indeed, we found that bile acid concentrations of cholic acid and deoxycholic acid, representing their reported abundances in the small and large intestines, significantly decreased pathogen growth, with a trend towards reduced growth in response to lithocholic acid (Figure 4C).

Other significantly altered metabolites included daidzein, an isoflavone derived from food sources such as soybeans and legumes, which has been reported to increase following vancomycin treatment^70^. Daidzein can be converted by gut microbes, such as *Lactobacillaceae*, to equol which shows strong antioxidant and anti-androgenic activity. Equol abundance was significantly decreased in abundance after vancomycin treatment, with a corresponding increase in abundance of daidzein, aligning with the loss of daidzein-metabolizing microbes such as *Lactobacillaceae* in the post-antibiotic gut environment (Figure 4A; Figure S4D)^71^. While equol did not impact *C. rodentium* growth, we found that daidzein significantly increased pathogen growth at 15-20 hours post-inoculation (Figure S4E). Many differentially abundant metabolites reflected expanded nutrient availability, including increased amino acid derivatives and oligo-and dipeptides, as well as simple sugars such as heptose monosaccharides (Figure S4F). These metabolites corresponded to the downregulation of several genes in the transcript module MEgreen related to the processing of sugars. For example, we observed a downregulation of genes related to the metabolism of galactonate, the oxidized form of galactose, such as *gntK* (encoding a gluconokinase), *dgoR* (encoding a D-galactonate utilization transcriptional regulator), *dgoD* (a galactonate dehydrogenase), *edd* (a phosphogluconate dehydratase), and *gntU* (a gluconate transporter). Instead, we observed an increase in expression of sugar transporter and carbohydrate porin-related genes (Figure 4D), suggesting direct environmental uptake.

### Cross-feeding between *Akkermansia* and *Citrobacter* at the mucus layer may promote early pathogen expansion

D-galactose and other simple sugars (L-fucose, N-acetyl-D-galactosamine [N-Gal], N-acetyl-D-glucosamine [NAG]) can be derived from host mucins. Given the increased abundance of *A. muciniphila*, a mucus-degrading bacterium capable of liberating simple sugars from the mucus layer^68^, we hypothesized that *C. rodentium* could benefit from cross-feeding at the mucus layer. Some *Bacteroides* species, such as *B. thetaiotaomicron*, are also mucus-dwelling bacteria capable of degrading mucin *O*-linked glycans^72,73^. To test whether mucus-degrading bacteria facilitate pathogen expansion, we inoculated mucus-producing HT29-MTX cells with *C. rodentium*, with or without live *A. muciniphila* under micro-aerophilic conditions (5 % CO_2_). In the presence of *A. muciniphila*, we observed significantly increased *C. rodentium* epithelial cell attachment, aligning with previous studies showing that mucus layer depletion by *A. muciniphila* allows for epithelial cell access by *C. rodentium*, as suggested in the aforementioned study (Figure 4E)^68^. However, we also noted increased planktonic *C. rodentium* in the presence of *A. muciniphila* (Figure S4G) suggesting increased overall pathogen growth.

While *C. rodentium* is capable of metabolizing mucin^74^ and growing in the presence of mucin as the sole carbon source, we hypothesized that the release of mucin-derived sugars by *A. muciniphila* provides a nutritive advantage during early pathogen colonization. We first grew *C. rodentium* in co-culture with one of two strains of *A. muciniphila* in mucin-rich media: the commercially available *A. muciniphila* ATCC strain or the human isolate *A. muciniphila* CC51001 Hb. We found that *A. muciniphila* increased *C. rodentium* growth after 24 hours in co-culture (Figure 4F). To determine whether this was a true cross-feeding relationship, we cultured *C. rodentium* in either fresh mucin-rich media or mucin-rich media pre-processed by *A. muciniphila* culture. *C. rodentium* was also cultured in media supplemented with water as a starvation control. Indeed, we found that *C. rodentium* grew to a higher bacterial load in *A. muciniphila*-processed mucin-rich media (Figure 4G). Together, these results indicate that the presence of *A. muciniphila* not only reduces the host mucus layer, allowing closer contact with epithelial cells, but also provides access to mucin-derived proteins and sugars, thereby allowing *C. rodentium* to cheat its way to population expansion.

We noted that *A. muciniphila* cross-feeding was beneficial only under anaerobic conditions, but not under 5% CO_2_, despite still observing viable *A. muciniphila* (Figure S4I). As the gut is known to oxygenate during infection, this relationship with *A. muciniphila* may provide an advantage during initial gut seeding^75^. Indeed, in our study and several others, the relative abundance of *A. muciniphila* and *B. thetaiotaomicron* has been shown to decrease post-*C. rodentium* infection, even in vancomycin pre-treated mice (Figure S4J). This may suggest that *A. muciniphila* and *B. thetaiotaomicron* provide a temporal advantage to initial pathogen establishment during early infection, to be later displaced by the expanding pathogen and changing intestinal architecture.

### Vancomycin widens colonization bottlenecks, allows for a large mucosal founding population size across the gut, and suggests low intra-species competition

Early infection events are critical to pathogen success. Under typical infection conditions extreme bottlenecks lead to significantly reduced pathogen diversity within the first 2 days of infection^22^. Considering the altered patterns of host colonization and overall infection success we hypothesized that natural bottleneck events, which protect the host from pathogen invasion, are compromised by vancomycin pre-treatment. To quantify bottleneck events within the gut, we used a previously described library of barcoded *C. rodentium*^22^. The library, which contains over 2,000 uniquely tagged *C. rodentium*, was created by inserting 30 random nucleotides into the pseudogene *flgN*, and validated for fitness as compared to the untagged wild-type strain^22^. Infecting mice with this tagged library allowed us to relate pathogen burden to genetic information via sequencing of colonized tissues and characterization of the tagged population (Figure 5A).

**Figure 5.**
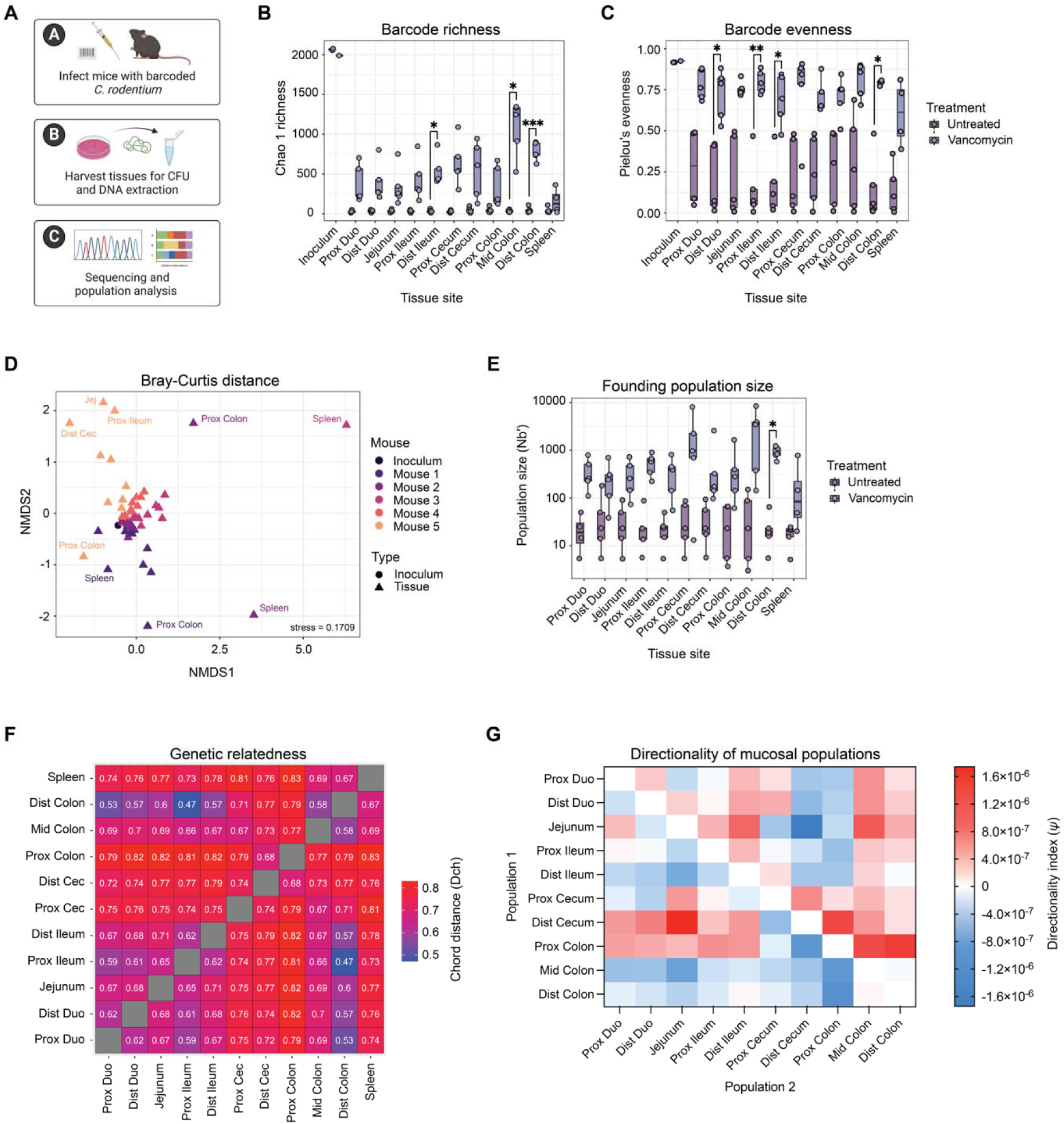
Barcoding reveals altered diversity and founding population sizes of tagged *C. rodentium* lineages in response to vancomycin pre-treatment. A) Overview of experiments using barcoded *C. rodentium* to explore population dynamics. In brief, a library of >2,000 tagged lineages was created by inserting a random 30-nucleotide barcode into a pseudogene of *C. rodentium*. The tagged library was then used to inoculate mice, harvesting infected tissues for sequencing and analysis of colonized lineages. B) Barcode richness (Chao1 index) across tissue sites at peak infection (day 7; N = 4-5). C) Barcode evenness (Pielou’s evenness) across tissue sites at peak infection (N = 4-5). Statistics represent a Mixed-effects model with Geisser-Greenhouse correction and Šidák’s multiple comparisons test. D) Non-metric multi-dimensional scaling (NMDS) analysis of sequenced inoculum and day 7 tissue samples in vancomycin pre-treated infected mice. Colours represent samples from individual mice. Triangles = tissue samples. Circles = inoculum samples. E) Founding populations size at day 7 post-infection across ten gut regions and the spleen, in untreated and vancomycin pre-treated mice (N = 4-5). F) Heat map showing the mean chord distance (Dch) between all intestinal samples at peak infection. Values greater than 0.2 indicate that populations are distinct (N = 5). G) Directionality index (ψ) calculations between intestinal tissues at day 7 post-infection (population 1 – population 2), where negative (blue) values indicate that population 1 is closer to the input population compared to population 2, and positive (red) values indicate that population 1 is farther from the input population (N = 5).

To compare gut regions between conditions at similar levels of colonization we focused on peak infection, collecting the mucosal (tissue-associated) population of colonizing *C. rodentium* across ten regional GI sites, as well as the spleen to represent systemic spread. We first investigated the diversity of barcode lineages present within the mouse gut. As expected, we found that barcode richness (Chao1) in the vancomycin group was significantly higher across all GI regions as compared to control animals, despite equal inoculum values, indicating increased diversity of colonizing lineages (Figure 5B). Interestingly, unlike in untreated animals where individual lineages came to dominate certain gut regions, in the vancomycin group lineages were more evenly distributed within the population (Figure 5C). Despite changes in alpha diversity of colonizing lineages within the gut, spleen pathogen richness was not altered as a result of antibiotic treatment (Figure 5B), likely because gut and systemic populations are subject to different bottlenecks. Exit of the GI tract to the systemic colonization is a major bottleneck during infection^22^, which may not be affected by antibiotic perturbation. However, we did observe an increase in Pielou’s evenness of barcode lineages in the spleen following vancomycin pre-treatment, indicating again a more evenly distributed systemic population. We further investigated beta diversity of barcode lineages by calculating the Bray-Curtis index. Compared to untreated infected animals, which showed stochastic colonization by individual mice (Figure S5C)^22,76^, the GI tissues of vancomycin pre-treated mice retain a composition similar to the inoculating population (Figure 5D). Exceptions included the spleen populations, indicating that the systemic bottleneck remains intact post-vancomycin treatment. Together, these data indicate a substantial increase in diversity of the colonizing pathogen population.

To quantify early colonization bottleneck size across the gut we estimated the founding population size (Nb′), a calculation that predicts the number of probable founders at any one site based on their relative frequencies within the population^47,77^. We found that Nb′ values were 10-fold higher across all gut regions in the vancomycin group compared to untreated controls (Figure 5E), with a significant increase in the distal colon region. This suggests that early bottleneck events are less severe after antibiotic perturbation. Interestingly, this removal of bottlenecks to regional colonization revealed additional patterns to Nb′ not seen in untreated infection. While average population size was the same across sites in untreated mice, vancomycin pre-treatment led to higher population size in the proximal cecum, mid-colon and distal colon regions, indicating that these sites are the least restrictive for pathogen colonization. Nb′ in the distal cecum, housing the cecal patch which is a preferred niche of *C*. *rodentium*, remained low. This is in keeping with the idea that the cecal patch region is competitive for initial competition while facilitating the expansion of colonizers to a high CFU burden^22^. While we saw an increase in systemic Nb′ as compared to untreated mice, spleen founding population size remained smaller than the Nb′ in the intestinal tissues, indicating that bottlenecks still remain to spleen colonization.

Considering the increased similarity of all tissue populations to the inoculating population and the high alpha diversity of colonizing barcoded *C. rodentium* lineages across the gut, we expected to find a high degree of relatedness between gut regional populations in vancomycin pre-treated mice. We investigated this by calculating the chord distance (Dch), a measure of the relatedness between populations. Small Dch values, below ∼0.2, indicate a high degree of relatedness between populations, whereas higher Dch values indicate that two populations are distinct. Interestingly, we found that composition across tissue sites was highly distinct (Figure 5F). This may indicate that gut sites are seeded independently of one another, and that regional pathogen populations are stable over time without being displaced by lumenal effluent. This was different from typical infection conditions in which we previously found that coprophagy was important for seeding of upper intestinal regions, leading to a high degree of relatedness between tissues^22^.

To further investigate the relationship between pathogen populations across intestinal sites, we next calculated the directionality index (ψ). ψ considers asymmetries in frequencies of tagged lineages, assuming that rare lineages are lost as population expansion occurs and abundant lineages will increase in frequency in the newly expanded population. ψ values will increase in the direction of expansion, such that a positive ψ value indicates that a population has likely expanded relative to the other, and a negative ψ value indicates the population is closer to the population origin. Results indicate that the mid-colon region had a negative ψ value compared to all other tissues, indicating that it may be the first site colonized in the vancomycin pre-treated gut, followed by the distal colon and distal ileum (Figure 5G). The cecum was unexpectedly one of the last sites predicted to colonize despite being the first in a typical infection setting, though this outcome could be skewed by severe bottlenecks in this region, making it the least representative of the inoculating population and thereby furthest from the population origin. We also noted evidence of possible retrograde colonization, in which each adjacent tissue section had a negative ψ value compared to the preceding section, with the exception of the distal colon. The spleen had a positive ψ value to all intestinal tissues indicating it is colonized after the gut, with the smallest distances to the upper small intestine and distal colon (Figure S5D), as is the case in untreated mice^22^. Taken together, our population analysis indicates major differences to regional mucosal colonization of vancomycin pre-treated mice.

We next sought to investigate the direct impact of the microbiota and metabolome on *C. rodentium* population size. By comparing pathogen population size within the colonic mucosa to the relative abundance of these commensal microbes at the time of infection, we identified a positive trend between pathogen Nb′ and *A. muciniphila* abundance (*p* = 0.08, rho = 0.9) and a negative trend to *B. thetaiotaomicron* abundance (*p* = 0.08, rho = –0.9; Figure S5E). We further repeated growth and attachment assays using the barcoded library (Figure S5F) and confirmed a significant increase in the founding population size of *C. rodentium* when grown in supernatants from cecal contents of vancomycin-treated mice as compared to untreated mice (Figure S5G). Similarly, we saw an increase in the founding population size of *C. rodentium* attached to HT29-MTX cells (Figure S5H). These data support our findings that the *Akkermansia*/*Bacteroides*-dominated microbiota reduces clonal competition and promotes the early, rapid expansion of *C. rodentium* within the host environment.

## Discussion

Antibiotic exposure is known to disrupt colonization resistance, yet we understand surprisingly little about how pathogens may exploit the antibiotic-exposed gut environment. We demonstrated that just two days of vancomycin exposure prior to pathogen challenge is sufficient to fundamentally alter infection dynamics, accelerating inflammation and increasing tissue pathology and disease severity (Figure 1).

Applying a barcoding approach allowed us to analyze pathogen population structure across intestinal sites and highlighted a critical consequence of vancomycin treatment–an increase in pathogen founding population size (up to 35-fold) and preservation of substantially greater genetic diversity within the infecting population (Figure 5). In contrast to untreated infections, where regional gut populations remain highly related^22^, vancomycin-treated mice exhibited reduced inter-site exchange (Dch ≍ 0.5-0.8), indicating that early-established lineages persist locally without displacement by clonal competitors. This resistance to clonal displacement likely stems from rapid, global gut colonization by day 2 post-infection, where in untreated animals, upper intestinal colonization occurs later and is shaped by coprophagy and lumenal flow^22^. Resistance to clonal displacement may directly limit therapeutic approaches that rely on competitive strain replacement^78,79^. In antibiotic-disrupted ecosystems where early founders persist and inter-lineages competition is relaxed, introduced competitors–whether commensal, attenuated, or engineered–may fail to displace established pathogen populations, reducing the efficacy of such interventions.

These findings align with growing evidence that mixed-strain and mixed-species infections are common in both acute and chronic disease, yet remain underappreciated in experimental models that rely on clonal inocula. In natural infections, which are inevitably non-clonal, relaxation of colonization bottlenecks may allow less competitive or poorly adapted lineages to establish and persist rather than being eliminated by host or microbial selection^80^. Indeed, whole-genome sequencing of *Enterococcus faecium* isolates from individual patients has revealed non-clonal populations harbouring variable resistance genes and plasmids^81^, while *Pseudomonas aeruginosa* infections frequently involve multiple strains whose competition accelerates antimicrobial resistance evolution^82^. By removing colonization bottlenecks, antibiotic exposure may therefore eliminate protective selective pressures that normally constrain pathogen diversity, facilitating co-colonization by mixed or poorly adapted strains, and potentially mixed species–an overlooked reality in clinical care.

Notably, increased disease severity did not correspond to heightened pathogen virulence. Instead, we observed reduced expression of LEE-encoded genes and decreased epithelial attachment in response to cecal metabolites from vancomycin-treated mice and to supernatants from *Bacteroides spp.* (Figures 2-3). This parallels previous findings that kanamycin treatment at peak *C. rodentium* infection induces a *Bacteroides* bloom associated with high fecal shedding but limited transmissibility^18^, consistent with our observation that *Bacteroides* suppresses *ler* expression. Although similar phenotypes were reported using a strain lacking the negative regulator *grlR*^18^, LEE expression is controlled by multiple transcriptional and translational regulators, allowing virulence expression to remain environmentally responsive. As *Bacteroides* are most dominant in the gut lumen, their detection by *C. rodentium* may act as a spatial cue that suppresses inappropriate T3SS expression away from the mucosa.

Rather than promoting virulence, post-antibiotic metabolic shifts appear to fuel rapid population expansion through nutrient abundance and loss of inhibitory metabolites such as bile acids, which normally constrain population density. Prolonged cefoperazone treatment similarly enriches carbohydrates and primary bile acids that promote germination of spore-forming *Clostridium difficile*^83^, suggesting a mechanism by which continued antibiotic treatment may inadvertently promote recurrent infection. This is particularly relevant given the widespread prophylactic use of antibiotics for recurrent infection-associated diseases, including urinary tract infections, where short courses are frequently administered in the absence of active infection^84^. Together, these data suggest that uncontrolled pathogen expansion can be as damaging to host health as hypervirulence.

Following vancomycin treatment, gut domination by *Akkermansia* and *Bacteroides* highlights their importance in shaping enteric infection. *A. muciniphila* blooms have been linked to increased *C. rodentium* disease severity, attributed to mucus thinning that facilitates epithelial access^68^. Under polysaccharide limitation, *A. muciniphila* upregulates mucin-degrading CAZymes to access mucus-derived glycans, eroding this protective barrier^68^. Here, we further identify a cross-feeding interaction between *A. muciniphila* and *C. rodentium*, adding a new dimension to this commensal-pathogen relationship (Figure 4). Although *B. thetaiotaomicron* did not enhance pathogen growth in our system, *Bacteroidales* spp. are known to promote *Enterobacteriaceae* expansion through nutrient exchange^85^, suggesting that *C. rodentium* may similarly *benefit in vivo*. These interactions reinforce the view that successful pathogens exploit, rather than disrupt, established microbial networks.

Together, our findings demonstrate that even brief antibiotic exposure can dismantle protective infection bottlenecks, enabling early, widespread colonization by genetically diverse pathogen populations. This competitive release shifts selection away from host adaptation and toward the persistence of costly or deleterious traits^80^. More broadly, our findings suggest that antibiotic-mediated relaxation of colonization bottlenecks may represent a general mechanism by which modern medical practices inadvertently favour pathogen diversity, persistence, and resistance evolution. Overall, our data reinforce that antibiotics act as ecological perturbations that extend far beyond pathogen killing, effectively reshaping microbial competition, metabolic constraints, and evolutionary trajectories.

## Acknowledgements

The authors would like to thank our colleagues in the Finlay laboratory for their support and assistance, especially L Thorson for their fundamental logistical support of the project. We thank A Lee for valuable input on RNA sequencing experiments and critical review of this manuscript. We would also like to thank H Bar-Yoseph for valuable consultation. This work was supported by grants from the Canadian Institutes of Health Research (CIHR) to BB Finlay (FDN-159935). SE Woodward received support for this project from a CIHR CGS-D Doctoral Scholarship, UBC Four Year Fellowship, and Dmitry Apel Memorial Scholarship. Supporting images were created with BioRender.com.

## Author Contributions

S.E.W. and S.L.V. conceived the project with guidance from B.B.F.. S.E.W., J.P.D., A.S.P., S.L.V., M.A.W., W.F., L.M.P.N., and J.F. performed experiments. S.E.W., A.S.P., and S.L.V. performed animal work. S.E.W., K.E.H., Z.K., and M.C. wrote bioinformatics pipelines and analyzed data. S.E.W. wrote the original draft of the manuscript with input from all authors. All authors revised the manuscript. B.B.F. acquired funding for the project and provided supervision.

## Data availability statement

The datasets generated during the current study are currently being deposited in the NCBI Sequence Read Archive (SRA). The accession numbers will be added once available. Meanwhile, data are available from the corresponding author upon reasonable request.

