## Supplementary material for "Vancomycin promotes key microbiota-pathogen interactions and removes protective bottlenecks to enteric infection": Table S1

Supplemental Figures and Tables

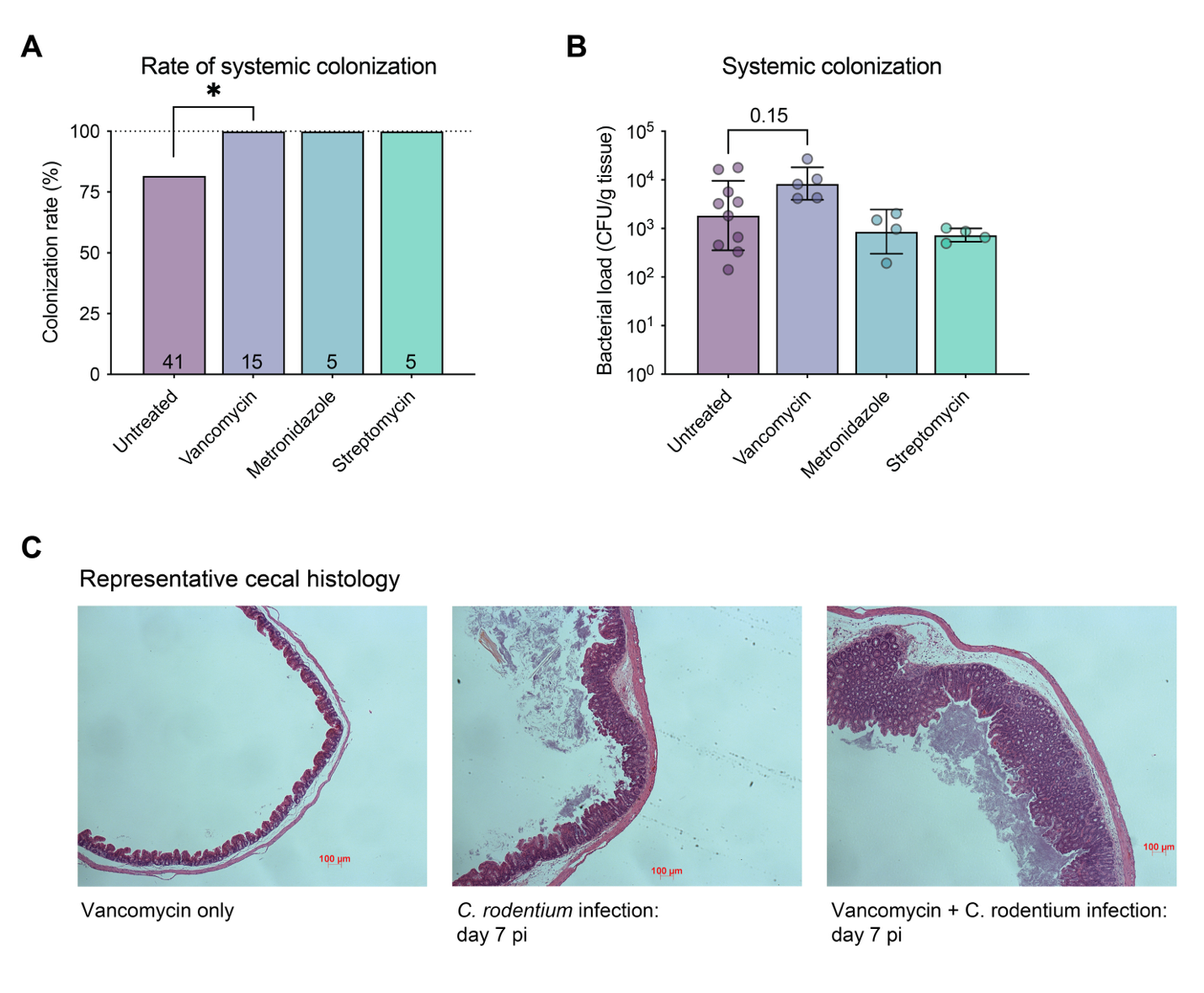

Figure S1. Effect of antibiotic treatment on systemic colonization by *C. rodentium*. A) Rate of systemic colonization, determined as the proportion of infected mice positive for CFU in the spleen at peak infection (day 7 post-infection). Statistics represent a Fisher’s exact test. B) Systemic pathogen burden at peak infection. Statistics represent a Kruskal-Wallis test with Dunn’s multiple comparisons test. C) Representative images of proximal cecum histology stained with hematoxylin and eosin. Sections shown of mice treated with vancomycin, infected with *C. rodentium*, and mice pretreated with vancomycin before *C. rodentium* infection. Infected tissues shown at day 7 post-infection (pi).

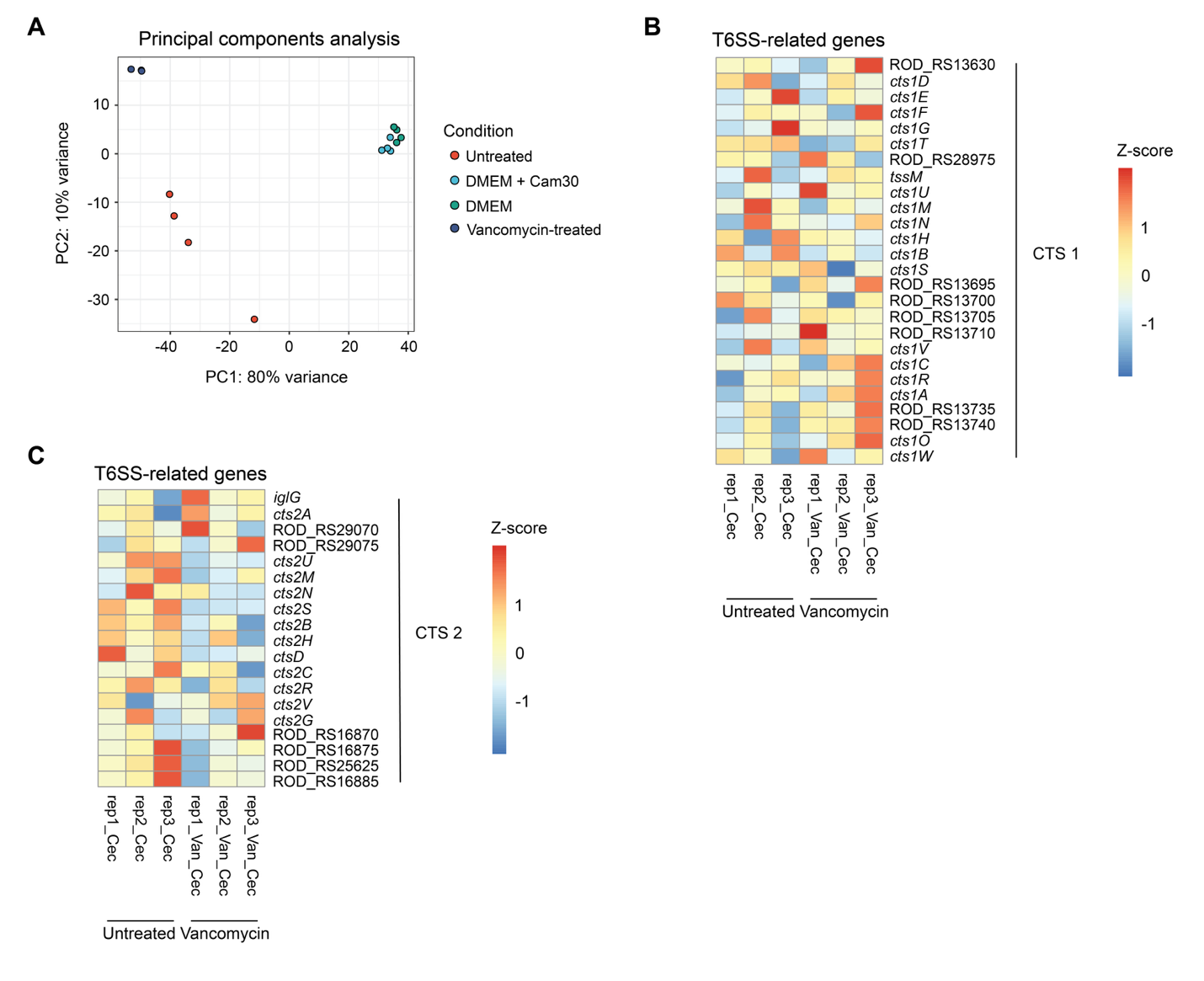

Figure S2. Expression of type VI secretion system genes in response to cecal supernatants from vancomycin pre-treated mice. A) Principal component analysis (PCA) of RNA sequencing normalized counts per condition, showing distinct clustering of *C. rodentium* in response to untreated versus vancomycin-treated cecal supernatants (N = 3-4). B) Heatmap showing Z-score expression of genes encoding type VI secretion system CTS-1 of *C. rodentium*. C) Heatmap showing Z-score expression of genes encoding type VI secretion system CTS-2 of *C. rodentium*. Vertical lines indicate the system (CTS-1 and CTS-2).

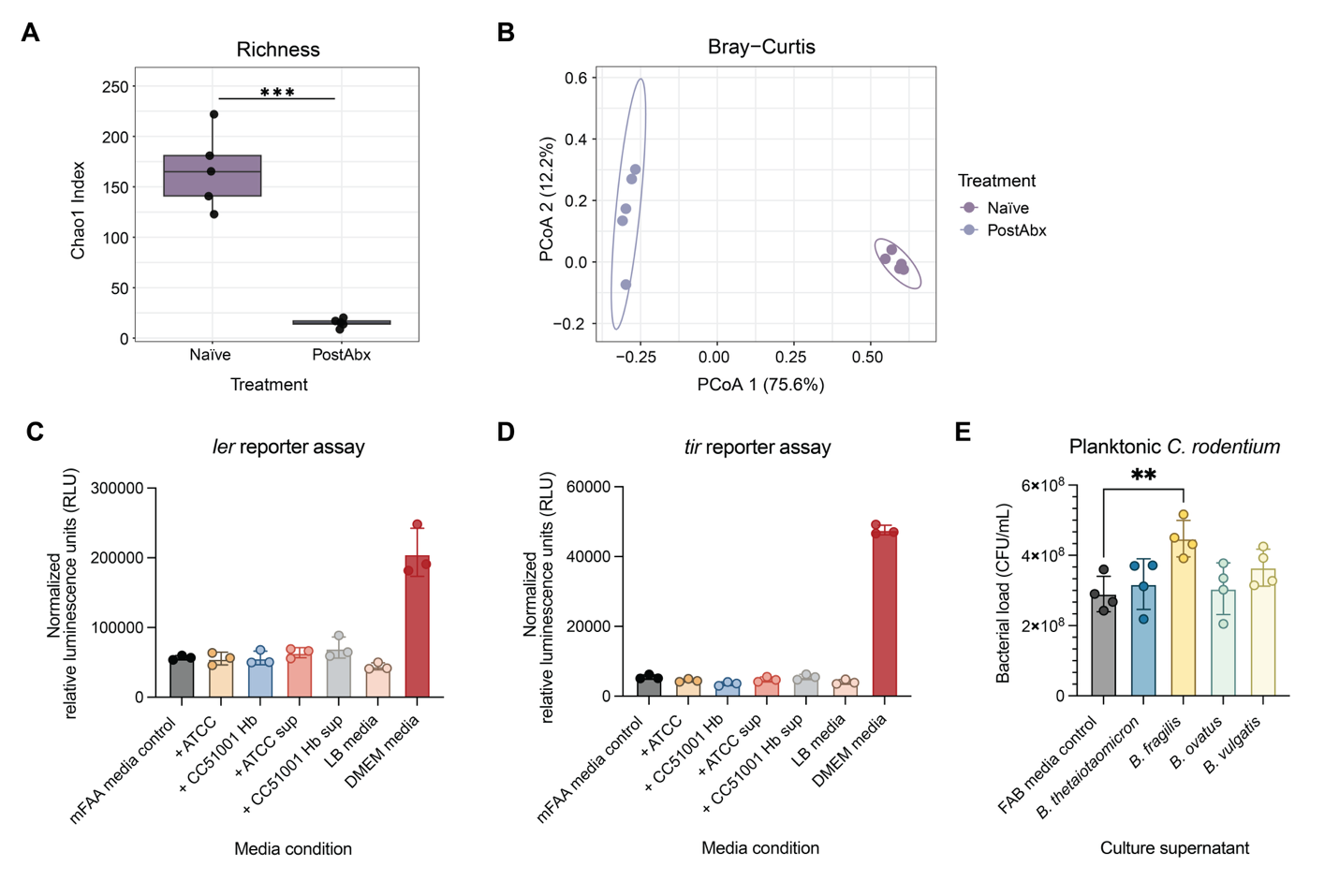

Figure S3. Effect of *Bacteroides* and *A. muciniphila* strains on *C. rodentium* growth and virulence. A) Chao1 richness of naïve mice and vancomycin-treated mice (N =5). B) Principal coordinate analysis (PCoA) of Bray-Curtis beta diversity index in naïve and vancomycin-treated mice (N = 5). C) *C. rodentium* *ler* gene expression in response to co-culture with live *A. muciniphila* or supernatants from *A. muciniphila* cultures, as measured by relative luminescence using a reporter strain. D) *C. rodentium* *tir* gene expression in response to co-culture with live *A. muciniphila* or supernatants from *A. muciniphila* cultures, as measured by relative luminescence using a *tir* reporter strain. Data normalized to *C. rodentium* CFU. E) Planktonic *C. rodentium* after 6-hour infection of HT29-MTX cells in response to *Bacteroides* culture supernatants.

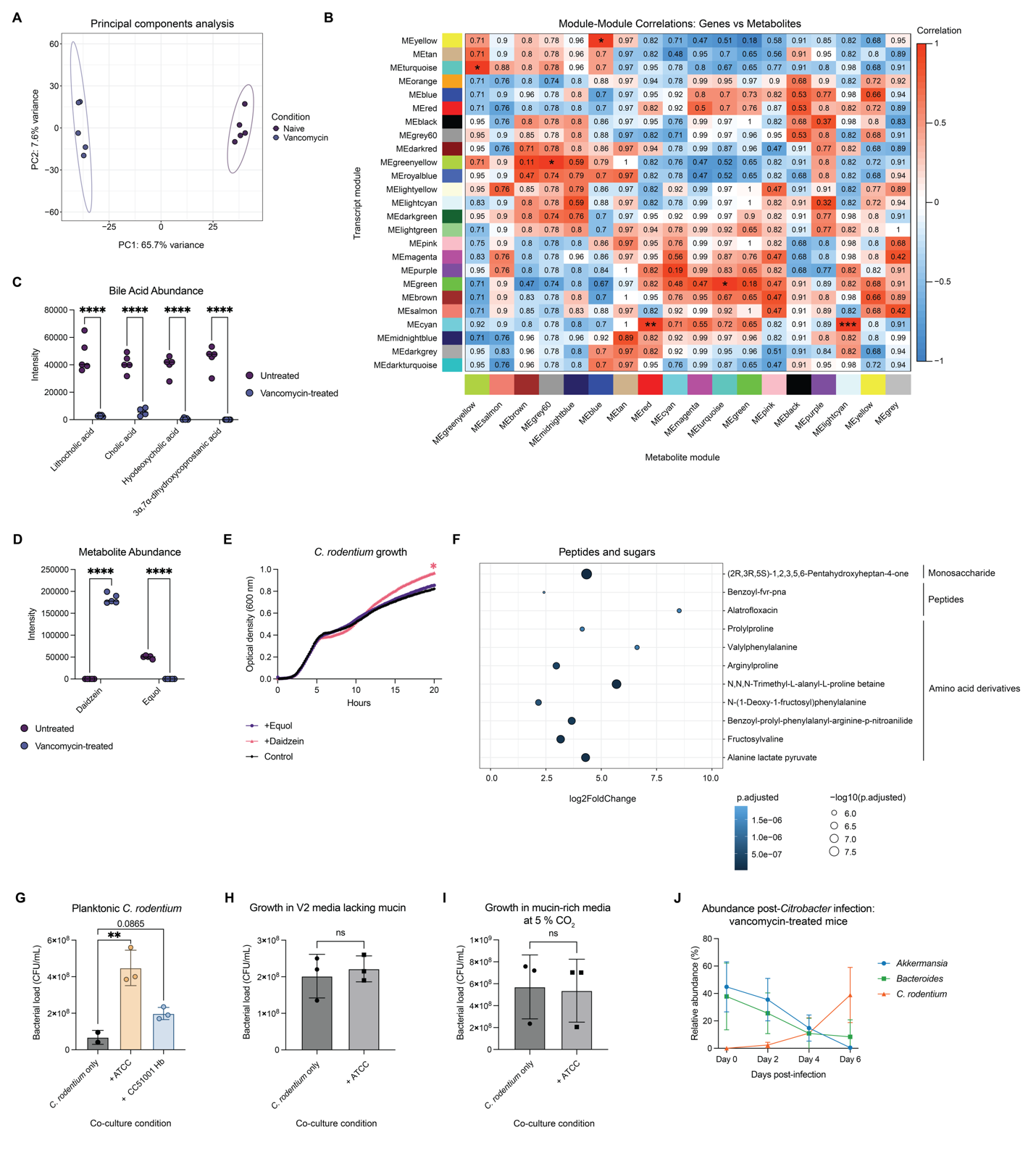

Figure S4. Transcript-metabolite relationships. A) Principal component analysis (PCA) of cecal metabolomic profiles, showing distinct clustering by pre-treatment condition (N = 5). B) Pearson correlation between modules assigned to *C. rodentium* transcripts and cecal metabolites using the WGCNA R package. Colours represent modules formed by transcripts or genes which correlate in abundance. C) Abundance of bile acid metabolites in the untreated and vancomycin-treated cecum as relative intensities (N = 5). D) Abundance of daidzein and equol in the untreated and vancomycin-treated cecum as relative intensities (N = 5). E) Growth of *C. rodentium* in LB media supplemented with equol or daidzein with OD 600 nm measurements taken at 10-minute intervals for 20 hours (N = 3 biological replicates, average of 3 technical replicates). F) Log2 fold change of amino acid derivatives, di- and oligo-peptides, and heptose monosaccharides in the vancomycin-treated cecal environment. G) Planktonic *C. rodentium* after 6-hour infection of HT29-MTX cells in the presence or absence of *Akkermansia* strains in co-culture. H) Growth of C. rodentium in V2 base media lacking mucin with or without the presence of *A. muciniphila* under anaerobic conditions (N = 3). I) Growth of *C. rodentium* in mucin-rich media with or without the presence of *A. muciniphila* under micro-aerophilic conditions (5% CO2; N = 3). J) Relative abundance of *A. muciniphila*, *B. thetaiotaomicron*, and *C.*

*rodentium* over time post-infection in vancomycin pre-treated mice (N = 5).

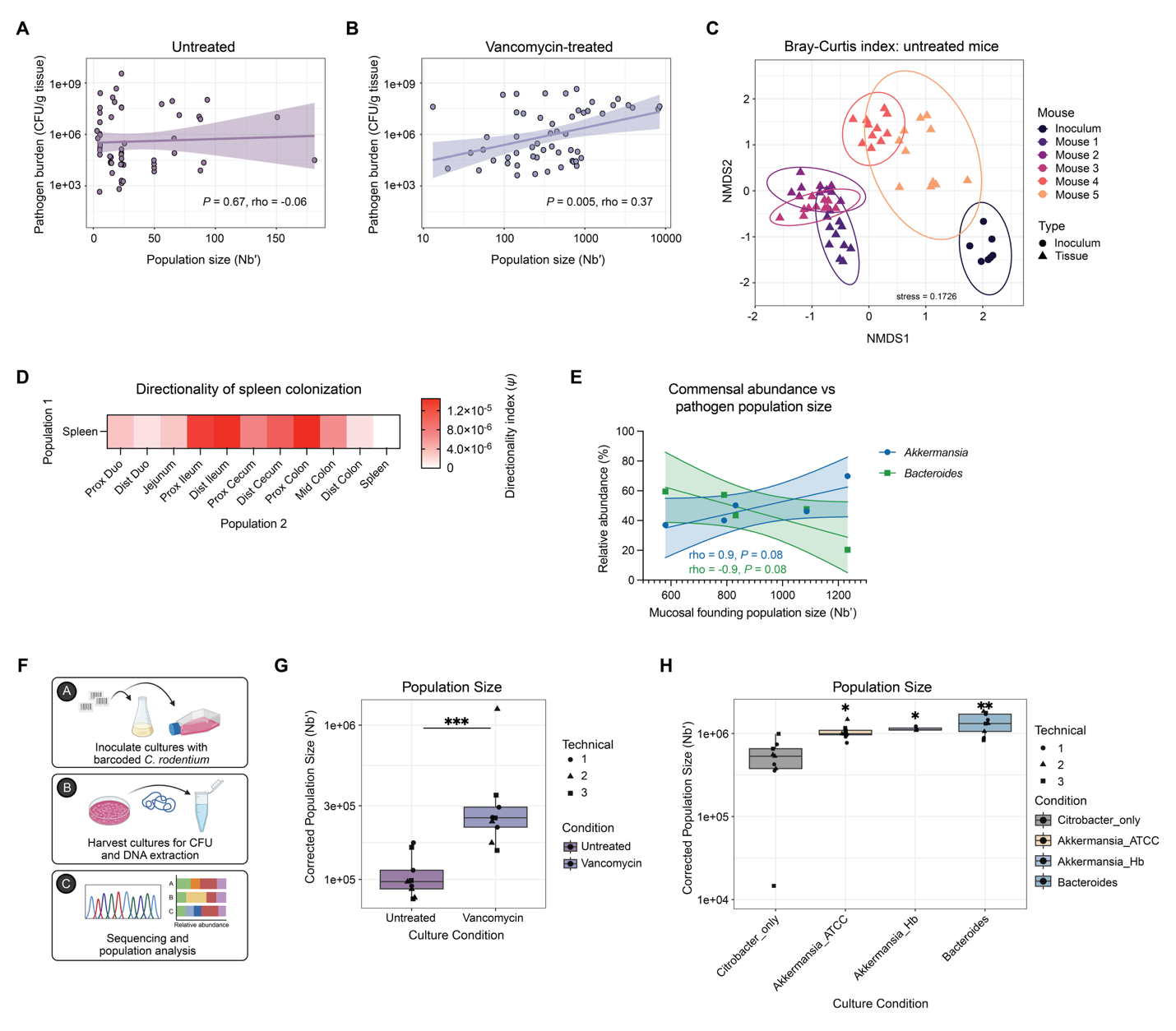

Figure S5. Analysis of barcoded lineages post-infection in untreated and vancomycin-treated mice. A) Spearman correlation of pathogen burden (CFU/g tissue) and founding population size (Nb') in untreated infected mice. Line represents linear regression +/- 95% confidence interval. *P*- and rho- values represent Spearman correlation values. B) Spearman correlation of pathogen burden (CFU/g tissue) and founding population size (Nb') in vancomycin pre-treated infected mice. Line represents linear regression +/- 95% confidence interval. *P-* and rho- values represent Spearman correlation values. C) Non-metric multi-dimensional scaling (NMDS) analysis of sequenced inoculum and day 7 samples in untreated mice. Colours represent samples from individual mice. Triangles = tissue samples. Circles = inoculum samples. D) Directionality index (*ψ*) calculations between the spleen and intestinal tissues at day 7 post-infection (population 1 – population 2), where positive (red) values indicate that population 1 is farther from the input population (N = 4). E) Spearman correlation of pathogen population size (Nb') with abundance of *A. muciniphila* and *B. thetaiotaomicron*. Line represents linear regression +/- 95% confidence interval. rho and *p*-values represent Spearman correlation values. F) Schematic of cell culture infection assays using the barcoded *C. rodentium* library. G) Corrected population size following *in vitro* growth of *C. rodentium* in cecal supernatants collected from untreated mice and vancomycin treated mice. (N = 3 biological replicates (each cultured in a pool of cecal content from 5 mice)). Statistics represent a Mann-Whitney test. H) Corrected population size of adherent *C. rodentium* following *in vitro* inoculation of HT29-MTX cells either in monoculture or in co-culture with *Akkermansia muciniphila* or *Bacteroides thetaiotaomicron* (N = 3 biological replicates). Statistics represent an Ordinary one-way ANOVA with Tukey’s multiple comparisons test performed on log-transformed values.

Table S1. Bacterial strains used in this study.

| **Strain** | **Description** | **Source or reference** |
| --- | --- | --- |
| DBS100 | Wild-type *C. rodentium*; ATCC 51459 | ^1^ |
| DBS100 Ler-lux | *ler* reporter strain made by inserting regulatory regions into pGEN-luxCDABE | This study. |
| DBS100 Tir-lux | *tir* reporter strain made by inserting regulatory regions into pGEN-luxCDABE | This study. |
| DBS100 *flgN*-barcoded library | 30-nucleotide barcode insertion in flagellar chaperone *flgN* with flanking constant regions inserted into the chromosome by single crossover; Cam^R^ | ^2^ |
| BAA-835 | *A. muciniphila* ATCC BAA-835 |  |
| CC51001 Hb | *A. muciniphila* human isolate | Dr. Emma Allen-Vercoe |
| *B. thetaiotaomicron* | ATCC 29148 |  |
| *B. fragilis* | Human fecal isolate | ^3^ |
| *B. ovatus* | Human fecal isolate | ^3^ |
| *B. vulgatus* | Human fecal isolate | ^3^ |

Table S2. Oligonucleotide primers used for 16S rRNA sequencing.

| Primer Name | Sequence | Description |
| --- | --- | --- |
| 515F | GTGCCAGCMGCCGCGGTAA | Forward primer to amplify 16S rRNA V4 |
| 806R | GGACTACHVHHHTWTCTAAT | Reverse primer to amplify 16S rRNA V4 |
